# Identification of immune-mitophagy related gene in PD based on single-cell sequencing and Mendelian randomization

**DOI:** 10.1101/2025.05.01.651665

**Authors:** Xiaolin Dong, Gang Wu, Lijuan Zhang, Qingyun Li, Yanping Li, Furong Jin, Rui Li, Yifan Ling, Yanming Xu

**Author notes:** Corresponding Author: Yanming Xu, 37 Guoxue Alley, Wuhou District, Chengdu, China.

## Abstract

Parkinson’s disease (PD) involves dysregulated mitophagy and immuneresponses, though the underlying mechanisms remain unclear. To identify key genes linking these pathways, we integrated single-cell RNA sequencing (GSE157783) with Mendelian randomization(MR) analysis, screening mitophagy-related genes (MRGs) and immune-related genes (IRGs). Differential expression analysis across PD cell subpopulations, combined with MR, revealed four causal genes: SLC11A1and DDX17(protective) and MRAS and PDIA3(risk). These genes were enriched in antigen presentation and calcium signaling pathways and exhibited dynamic expression in astrocytes and microglia during differentiation. Subsequent protein-protein interaction(PPI) network, regulatory (SCENIC), and drug-target analyses further characterized their roles. Validation in MPTP-induced PD mice confirmed behavioral deficits and altered expression of these genes, supporting their functional relevance. Our findings highlight SLC11A1, DDX17, MRAS, and PDIA3 as critical mitophagy-immune hubs in PD, offering mechanistic insights and therapeutic targets.

**Author Summary:** PD is a neurodegenerative disorder linked to mitochondrial dysfunction (mitophagy) and immune dysregulation, yet the interplay between these mechanisms remains poorly understood. Using integrative single-cell sequencing and MR analyses, we identified four key genes-SLC11A1, DDX17, MRAS, and PDIA3-that bridge immune and mitophagy pathways in PD pathogenesis. These genes exhibit causal relationships with PD risk, offering novel insights into diagnostic and therapeutic strategies.

## Introduction

Parkinson’s disease (PD) is the second most common neurodegenerative disease in the world, with main symptoms such as static tremor, motor delay, limb stiffness, and postural instability [1,2], seriously affecting the lives of patients. The incidence rate over 65 years old is 1.7% in China and is expected to rise to 5 million in 2030 [3,4]. However, the pathogenesis of PD has not been fully understood. Currently, it is believed that in addition to genetic susceptibility, environmental and neurological aging factors, such as oxidative stress, cell apoptosis, and mitochondrial dysfunction, lead to significant degeneration and loss of dopaminergic (DA) neurons [5]. Due to atypical non-motor symptoms (such as mood, sleep disturbances, visual impairments, fatigue, and so on) in the early stages, a high rate of misdiagnosis in early [6]. Therefore, studying the pathogenesis of PD and searching for molecular markers for early diagnosis and effective treatment are particularly important [7].

The loss of DA in the substantia nigra is the main pathology [8, 9]. Mitochondria, as the main energy generator in cells, make DA neurons extremely sensitive to them [10]. This also indicates that mitochondria play a crucial role in neuronal death. More importantly, as a regulatory mechanism for mitochondrial self-monitoring, mitophagy can clear abnormal mitochondria to ensure the quantity and quality of mitochondria. Previous studies have shown that if mitophagy were abnormal, high levels or prolonged damage to mitochondria can damage neurons and accelerate the progression of PD [11]. Mitophagy is also related to the immune system, involving the recognition and degradation of pathogens, antigen presentation, lymphocyte development, and regulation of inflammation [12,13]. A nationwide epidemiological study showed that the overall excess risk of PD in patients with autoimmune diseases is about 33% [14]. Additionally, immune function plays a complex role in PD while also demonstrating neuroprotective effects [15,16]. Although mitophagy and the immune system are considered key pathogenic mechanisms in PD, their interaction in PD has not been reported and requires further investigation.

Therefore, utilizing single-cell analysis and magnetic resonance imaging, our study seeks to identify key genes associated with MRGs and IRGs in PD, investigating their biological mechanisms and potential drug targets. Through this research, we aim to provide important insights that can help our understanding of how PD develops and identify potential new diagnostic genes for the disease.

## Results

### Astrocytes and Microglia were identified as candidate key cell subpopulations

A total of 39131 cells were obtained from GSE157783 through quality control (Figure S1a and S1b). After standard data processing, 2,000 highly variable genes were identified (Figure 1a). The first 30 PCs were selected for subsequent analysis according to the principal component inflection point diagram and gravel diagram (Figures 1b and 1c). Then, 27 different cell clusters were identified by UMAP clustering analysis (Figure 1d). Based on the marker gene and the Single R package, these cell clusters were annotated to 12 cell subpopulations, namely Astrocytes, CADPS2+ neurons, DaNs, Endothelial cells, Ependymal, Excitatory, GABA, Inhibitory, Microglia, Oligodendrocytes, OPCs and Percytes (Figure 2a and 2b). In addition, The proportion of Oligodendrocytes in PD and control samples was the highest, 46.83% and 59.32%, respectively (Figure 2c). Subsequently, Astrocytes and Microglia showed significant differences between the PD and the control groups, so they were used as candidate key cell subpopulations for follow-up analysis (Figure 2d).

**Figure 1.**
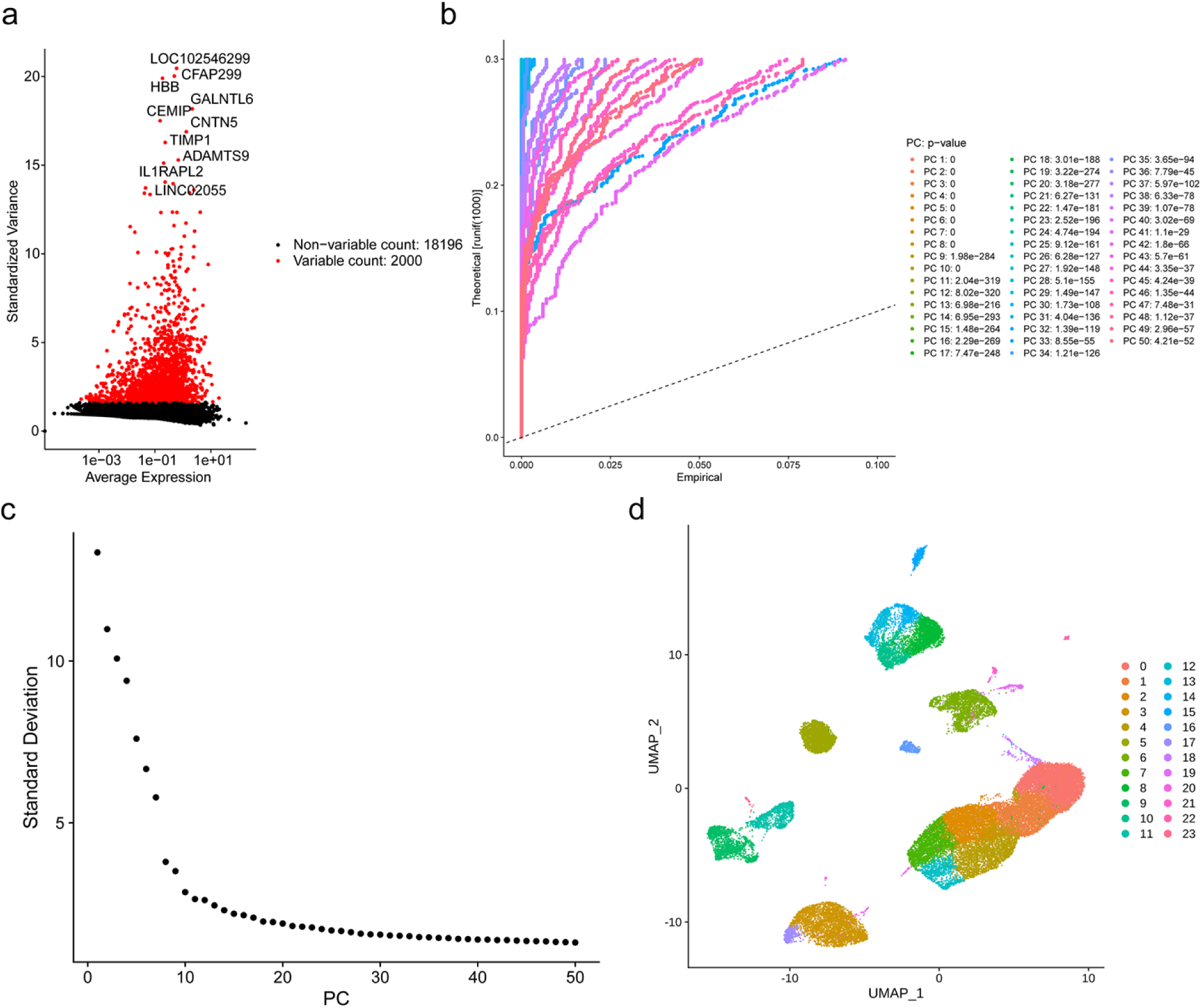
Single-cell atlas in GSE157783 dataset. (a) High variant gene screening. The horizontal axis is the average expression level, and the vertical axis is the standard variance. (b) PCA component analysis. (c) Graph of principal components. The location of the principal component 30 is already at a plateau stage. (d) UMAP diagram of dimensionality reduction clustering of single cell sequencing data.

**Figure 2.**
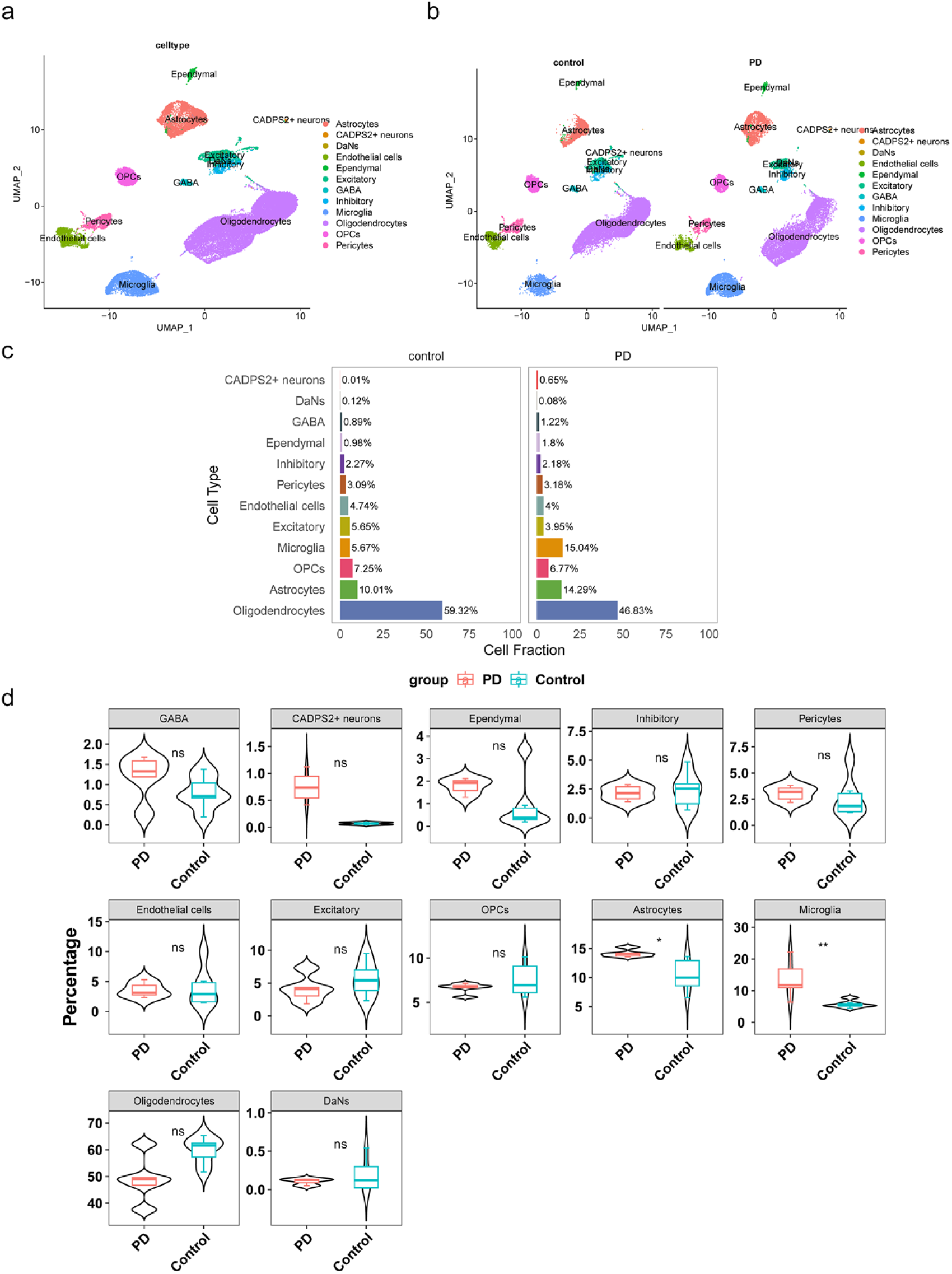
Identification of key cell clusters. (a-b) The UMAP plot of 12 cell clusters from 30 principal components. Cells from different clusters are marked by colors. (c-d) The cell fraction of PD vs Control.

### Cell communication of candidate key cell subpopulations

In order to search for cellular communication between candidate key cell subpopulations and other cell subpopulations, network maps, and heat maps were drawn. There were strong interactions existed between Astrocytes and Ependymal, Excitatory, and Inhibitory. There were strong interactions between microglia and OPCs, as well as astrocytes, endothelial cells, and ependymal cells (Figure 3a and 3c, Figure S2a). In addition, ligand-receptor interactions could also be observed. For instance, Astrocytes and DaNs communicated with each other through TGFB2 (ligand)-ACVR1B_TGFbR2 (receptor) (Figure S2b, Table S1). Furthermore, GSVA was carried out to explore the signaling pathways for differences between two candidate key cell subpopulations. The results showed that there were differences in the enrichment pathways of two candidate key cell subpopulations, such as tnfasignalingvianfkb, hypoxia, and cholesterol homeostasis (Figure 3d).

**Figure 3.**
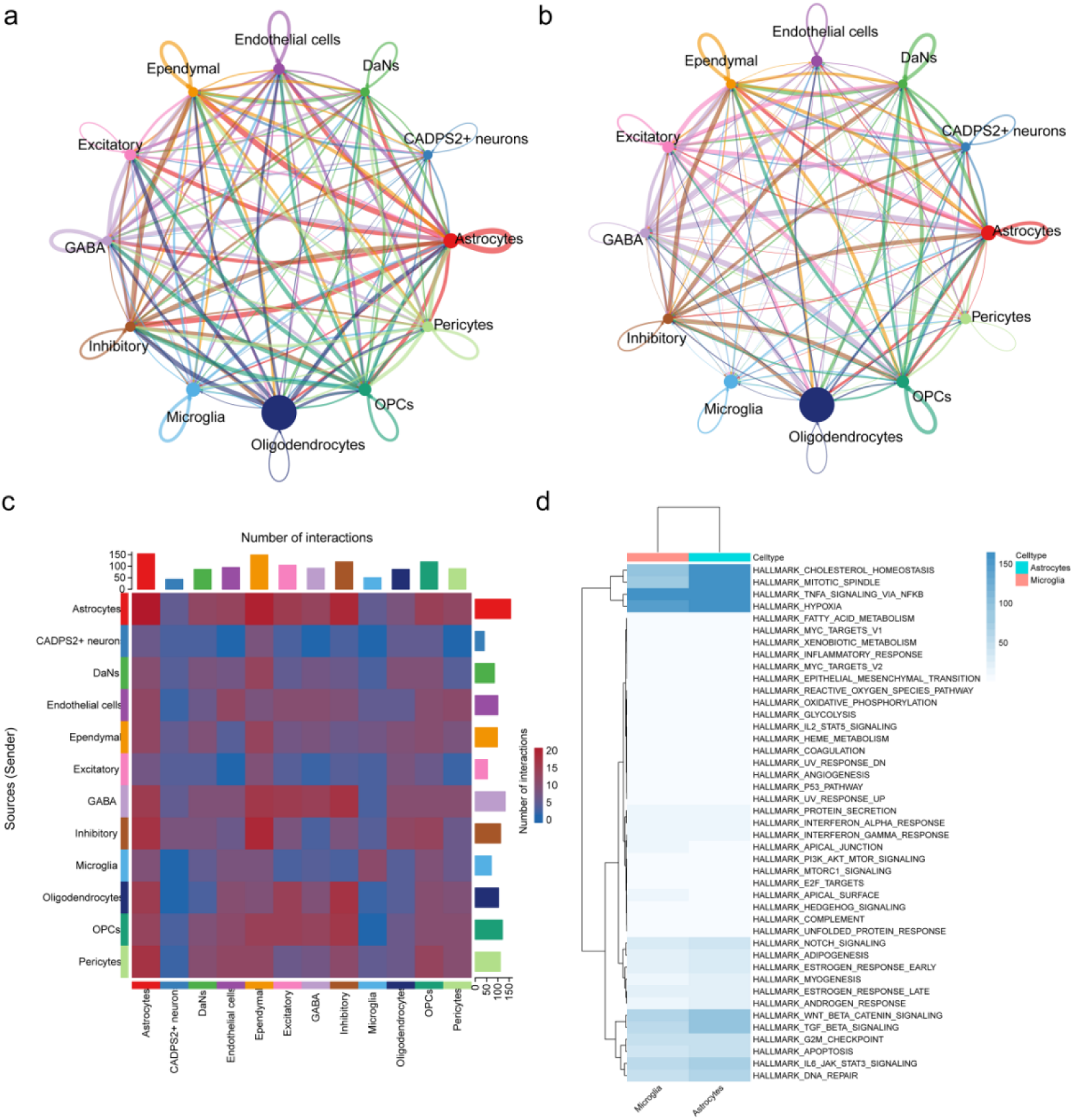
Cell communication. (a-b) Cellular communication between the different cell types. Arrows indicate the direction, and the line thickness indicates the number of interactions between cells (left) or weighted (right). (c) Heat map of number of interactions. (d) GSVA of the key cells, and the enrichment analysis. The white blue gradient indicates the change of-log10.

### The 58 candidate key genes were enriched in the cytokine−mediated signaling pathway

Based on differential expression analysis, a total of 526 DEGs were identified in Microglia, with 287 upregulated and 239 downregulated genes (Figure 4a). In Astrocytes, 878 DEGs were identified, with 475 upregulated and 403 downregulated genes (Fig 4b). The union of DEGs from the two cell clusters resulted in 1282 unique DEGs after removing duplicates. Further correlation analysis involved selecting genes with a correlation greater than 0.7 (|cor|>0.7) and p<0.01 between IRGs and MRGs for subsequent analysis. Specifically, 682 and 685 genes were respectively selected from Astrocytes and Microglia (Table S2-S3). Upon taking the intersection of the results, a total of 453 IRGs-MRGs were obtained (Table S4). The 58 candidate key genes were obtained by intersecting 1282 shared DEGs and 453 IRGs-MRGs (Figure 4c). An enrichment analysis was performed to explore the biological functions of candidate key genes. In GO items, candidate key genes were mainly enriched in cytokine−mediated signaling pathways, pattern recognition receptor signaling pathway ext, intrinsic apoptotic signaling pathway, etc. (Figure 4d). The result of KEGG showed that candidate key genes were involved in the IL-17 signaling pathway, MAPK signaling pathway TNF, signaling pathway, etc (Figure 4e).

**Figure 4.**
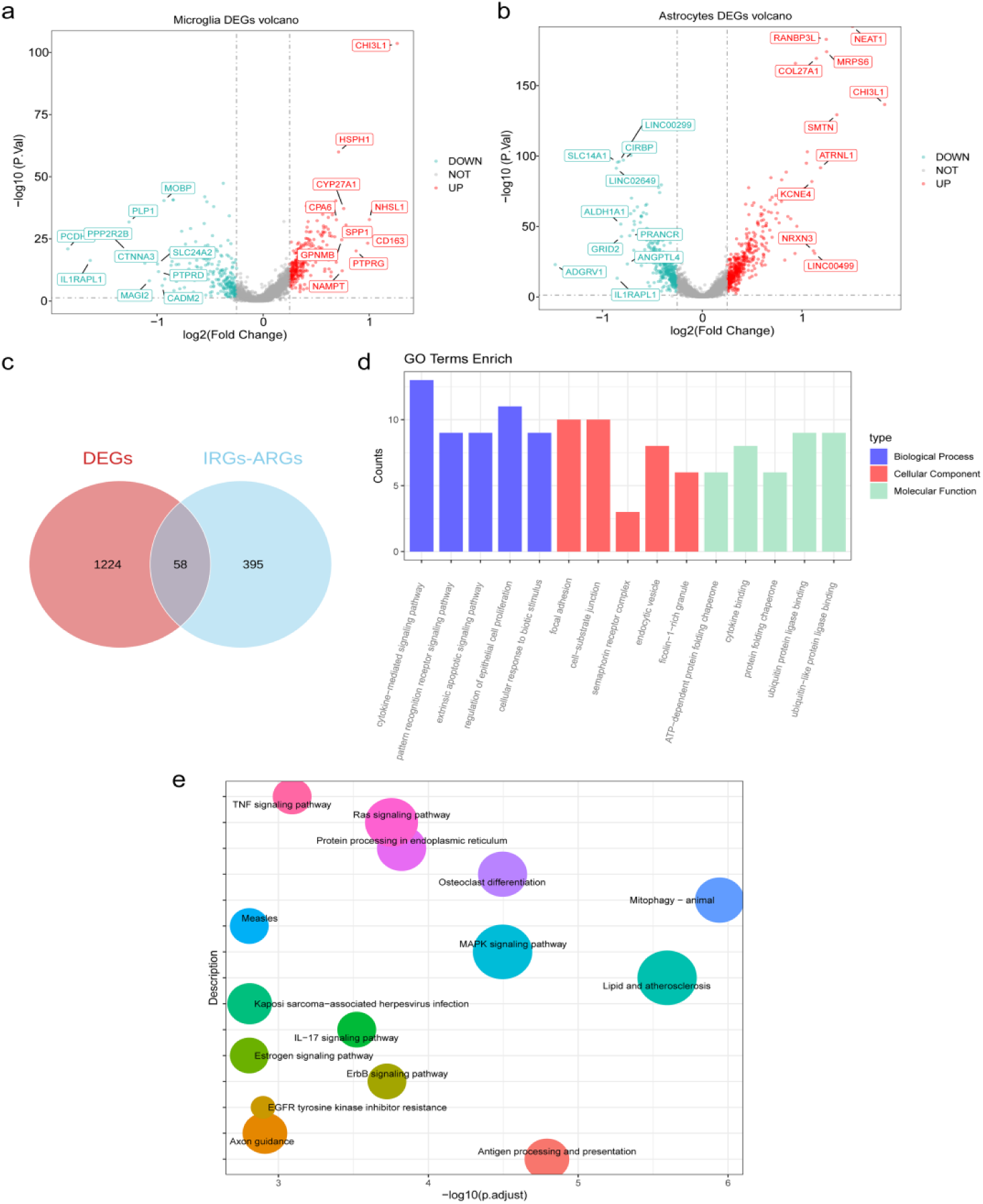
Identification of DEGs in GSE157783 dataset. (a) Volcano map of differentially expressed genes between sample groups. PD VS Control, the horizontal coordinate difference multiple (logarithm of 2), and the vertical coordinate represents-log10 (p. adj), Each dot represents a gene. (b) Differential expression gene density heat map. Red indicates high expression and blue indicates low expression. (c) The Veen graph of the intersection of DEGs and IRGs-MRGs. (d) GO enrichment chords. Different colors distinguish different pathways, and the genes pointed by the strings are the genes enriched in this pathway. (e) KEGG enrichment analysis tree of candidate key genes. The grid size represents the number of genes enriched in that term.

### There is a significant causal relationship between SLC11A1, DDX17, MRAS and PDIA3 and PD

After screening for IVs, 35 candidate key genes were retained for MR analysis, among which four genes had a significant causal relationship with PD in the IVW method of UVMR analysis (Table S5). SLC11a1 related dataset (equal-a-ENSG00000018280) 17,576 SNPs from 31,684 samples. DDX17 related dataset (equal-a-ENSG00000100201) 18,143 SNPs from 31,470 samples. MRAS-related dataset (equal-a-ENSG00000158186) has 16,813 SNPs from 31,684 samples. PDIA3-related dataset (equal-a-ENSG00000167004) 14,760 SNPs from 26,609 samples. After filtration, The numbers of SNPs associated with SLC11a1, DDX17, MRAS, and PDIA3 are 21, 10, 22, and 10, respectively.

In addition, SLC11a1 (odds ratio (OR) = 0.95, *p* value= 0.022), DDX17 (OR = 0.92, *p* value = 0)were protection factors for PD, while MRAS(OR = 1.25, p value = 0) and PDIA3 (OR = 1.11, *p* value = 0.037) were risk factors for PD. The slope in the scatter plot was negative, and the MR effect value of the forest plot was less than 0, which further supported SLC11a1 and DDX17 reduced the risk of PD, while MRAS and PDIA3 were the opposite (Figure 5a and 5b). In addition, the distribution of SNPs in the funnel plot was basically symmetrical and homogeneous on the whole, indicating that the MR analysis conformed to Mendel’s second law(Figure 5c). A sensitivity analysis was performed to assess the reliability of the MR analysis. The *p*-value of 4 genes was greater than 0.05, indicating there was no heterogeneity between the exposure and outcome datasets (Table 3). In addition, horizontal pleiotropy was not present in any of the four genes (*p*-value> 0.05) (Table 4). According to the results of the LOO sensitivity test, there were no extra sensitive IVs, indicating that MR analysis satisfied Mendel’s second law(Figure 5d). Subsequently, MVMR analysis was used to determine the influence of four genes on PD. The results showed that the four genes for PD were no longer significant, which may mean the effects of these genes on PD were interrelated to some extent (Table 5, Figure 5e). In summary, SLC11a1, DDX17, MRAS, and PDIA3 were identified as key genes with a significant causal relationship with PD.

**Figure 5.**
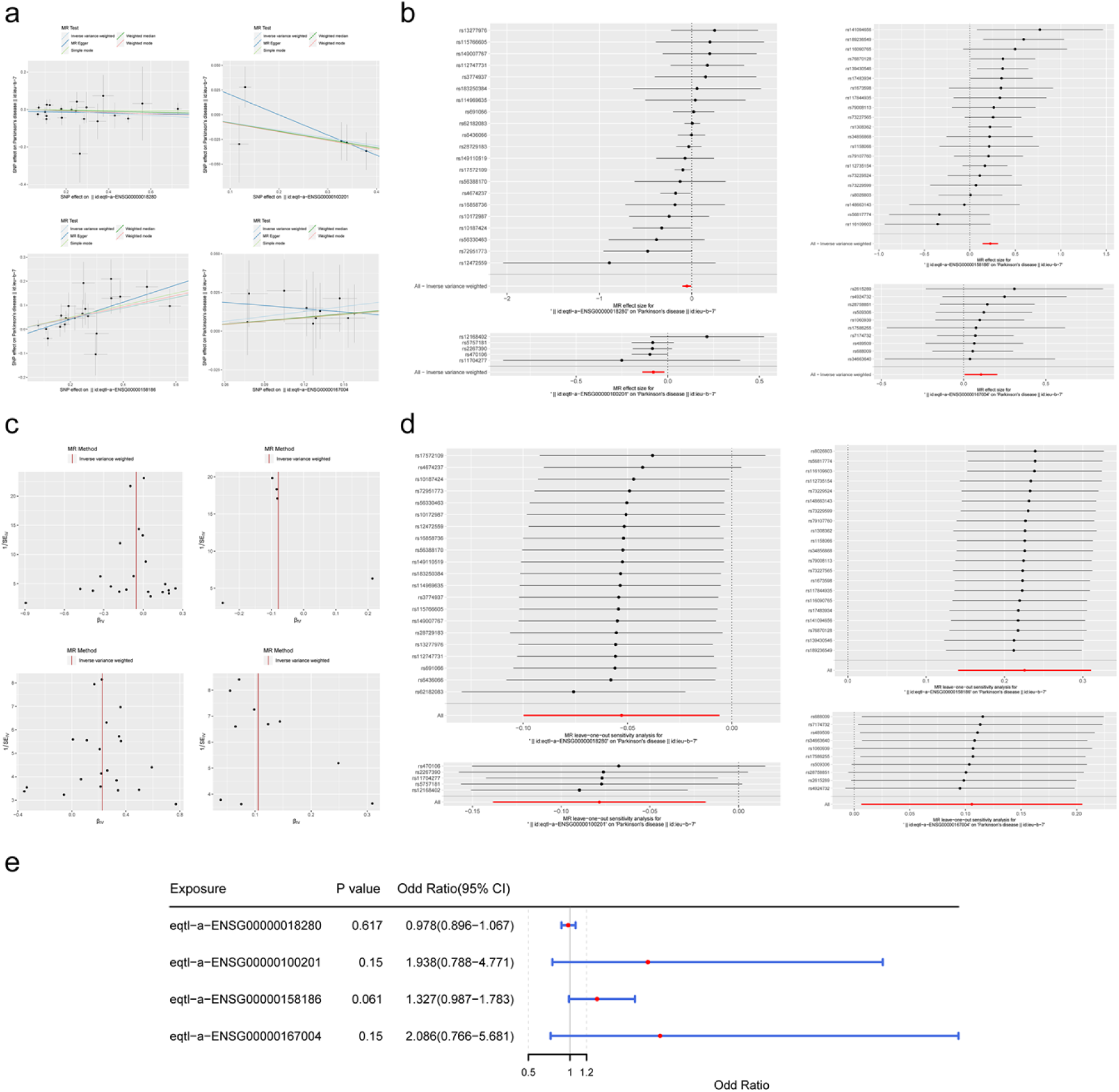
MR analysis. (a) Correlation analysis and the scatter plot of SLC11a1, DDX17, MRAS and PDIA3. (b) Forest plot of the diagnostic efficacy of SLC11a1, DDX17, MRAS and PDIA3. (c) Funnel diagram of SLC11a1, DDX17, MRAS and PDIA3. The IVs are symmetrically distributed in the funnel plot, then indicating that SNPs in this study follow the Mendelian Model’s second law random distribution rule. (d) LOO sensitivity test of SLC11a1, DDX17, MRAS and PDIA3. All points are on the right side of 0, and there is no significant shift in the distribution, indicating that no exceptionally sensitive instrumental variables exist. (e) Multivariate MR forest plot.

### Differential expression analysis of key genes and construction of nomogram

In the dataset GSE22491, the Wilcoxon rank-sum test was utilized to conduct a differential analysis of the expression levels of key genes. The results revealed significant differences (P<0.05) in the expression of SLC11a1, DDX17, and PDIA3 between the PD group and the control group (Figure 6a). This study constructed a nomogram model based on the expression levels of four genes, DDX17, MRAS, PDIA3, and SLC11A1, in the training set GSE22491 (Figure 6b). The calibration curve with a p-value > 0.05 indicated that the nomogram prediction model had a certain degree of accuracy in predicting the probability of PD (Figure 6c). The results of the DCA curve suggested that the nomogram possesses considerable predictive accuracy (Figure 6d). The AUC value of the ROC curve being greater than 0.7 further demonstrated the good predictive accuracy of the nomogram (Figure 6e).

**Figure 6.**
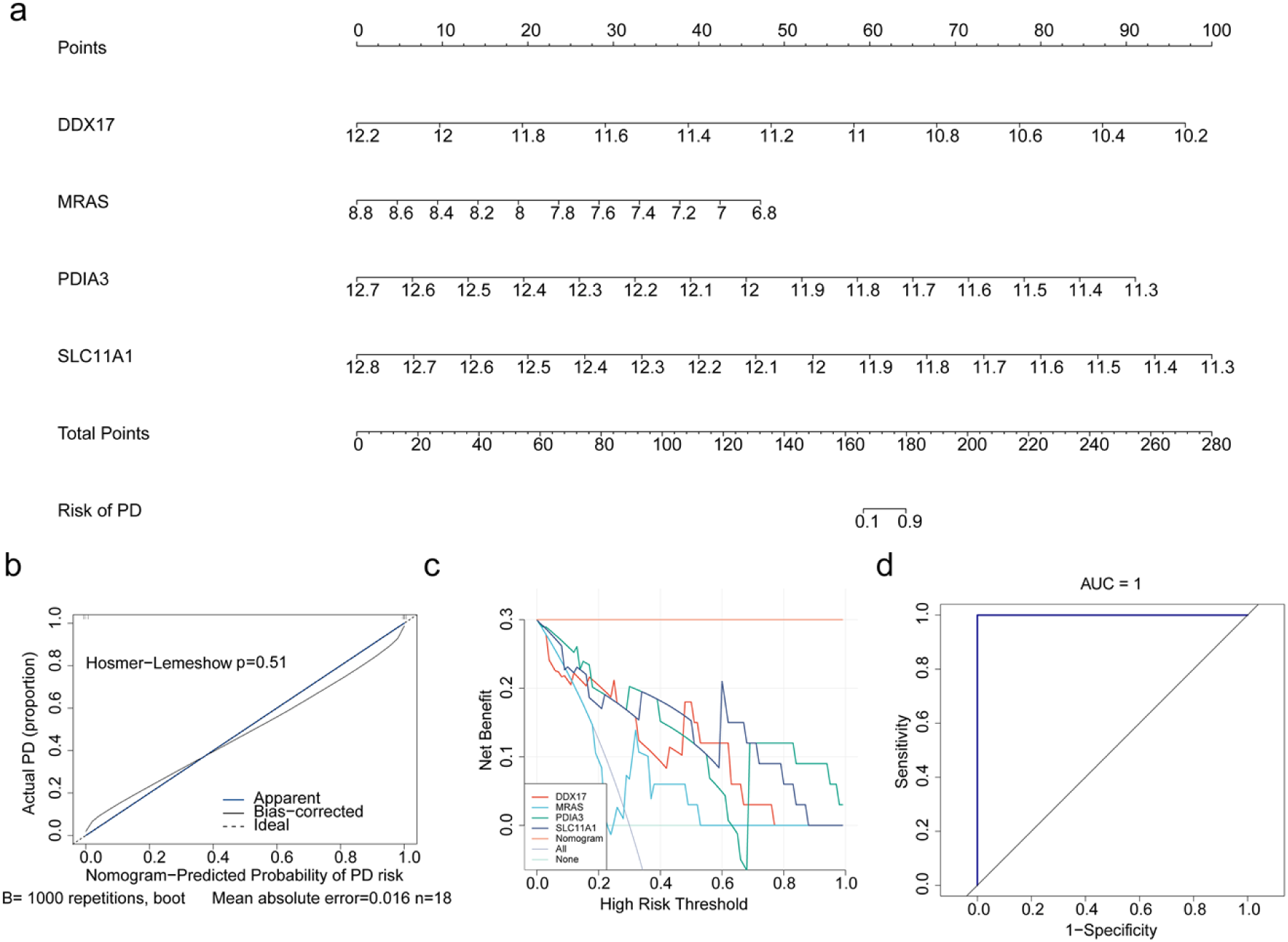
Differential expression analysis of key genes and construction of nomogram. (a) Differential expression of key genes in the training set GSE22491. *\*: P*<0.05; *\*\**: *P*<0.01; ns: no significance. (b) Construction of nomogram. (c) The calibration curve of the nomogram. (d) The decision curve analysis of the nomogram. (e) The receiver operating characteristic (ROC) curve of the nomogram.

### Most of the key genes were related to antigen processing and presentation

In order to understand the pathway where key genes were enriched, GSEA was performed. The result of GSEA found that SLC11a1, MRAS and PDIA3 were all enriched in antigen processing and presentation, and DDX17 and MRAS were all involved in calcium signaling pathway (Figure 7a-d). In addition, a strong interaction between SLC11a1 and SLC11a2 was found in the PPI network (Figure 7e).

**Figure 7.**
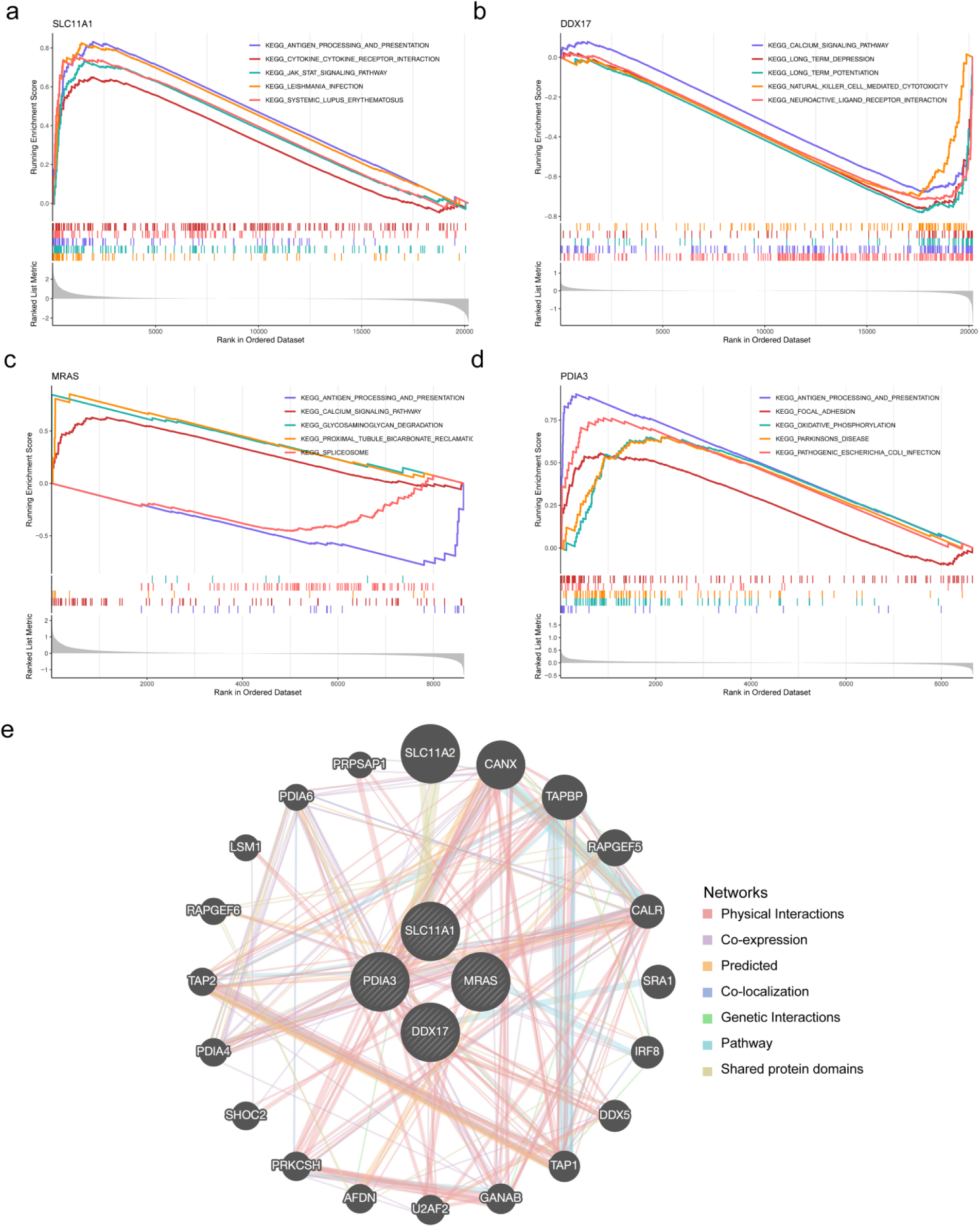
GSEA enrichment results of four biomarkers. The ordinate represents the enrichment fraction, and enrichment score (ES) is positive, indicating that a certain functional gene set is enriched in front of the sequencing sequence, indicating that it is positively correlated with gene enrichment. ES is negative, indicating that a certain functional gene set is enriched behind the sequencing sequence, indicating that it is negatively correlated with gene enrichment. The horizontal coordinates represent genes, and each little vertical line represents a gene. (a) SLC11a1. (b) DDX17. (c) MRAS. (d) PDIA3. (e) Protein co-expression network of four biomarkers.

### Four key genes played a role in the development of key cell subpopulations

The expressions of DDX17, MRAS, and PDIA3 in Astrocytes were significantly up-regulated in the PD group, while SLC11a1 was not significant. In Microglia, the expression of PDIA3 and MRAS were significantly up-regulated in the PD group, while SLC11a1 was the opposite, with DDX17 not exhibiting significant changes. Based on the differences in the expression of key genes between PD and control samples, we further identified Astrocytes and Microglia as a key cell subpopulation (Figures 8a and 8b). Both cells were found to be up-regulated in the PD group. Subsequently, the key cell subpopulations were further clustered based on marker genes. Astrocytes and Microglia were divided into 9 and 11 new subtypes, respectively (Figure 8c and 8d). Then, the nine astrocyte clusters were classified as nine astrocyte subtypes, which were astrocyte FOXG1, astrocyte SHISA6, astrocyte LGR6, astrocyte ZNF536, astrocyte GBP2, astrocyte RSPO2, astrocyte ROBO2, astrocyte ITGAX, and astrocyte IFI27L2; the 11 microglia clusters were classified as 10 MG subtypes, which were MG IL7R, MG IL1RAPL1, MG EGFR, MG LYVE1, MG FOS, MG CACNA1A, MG CCDC26, MG TMEM163, MG MKI67 and MG GPNMB (Figure 8e and 8f). Moreover, the expression patterns of key genes showed that DXX17 was expressed in each subtype of astrocytes and MG (Figure 8g and 8h). These results suggested that DDX17 might influence the subtypes of astrocytes and MGs involved in the development of PD.

**Figure 8.**
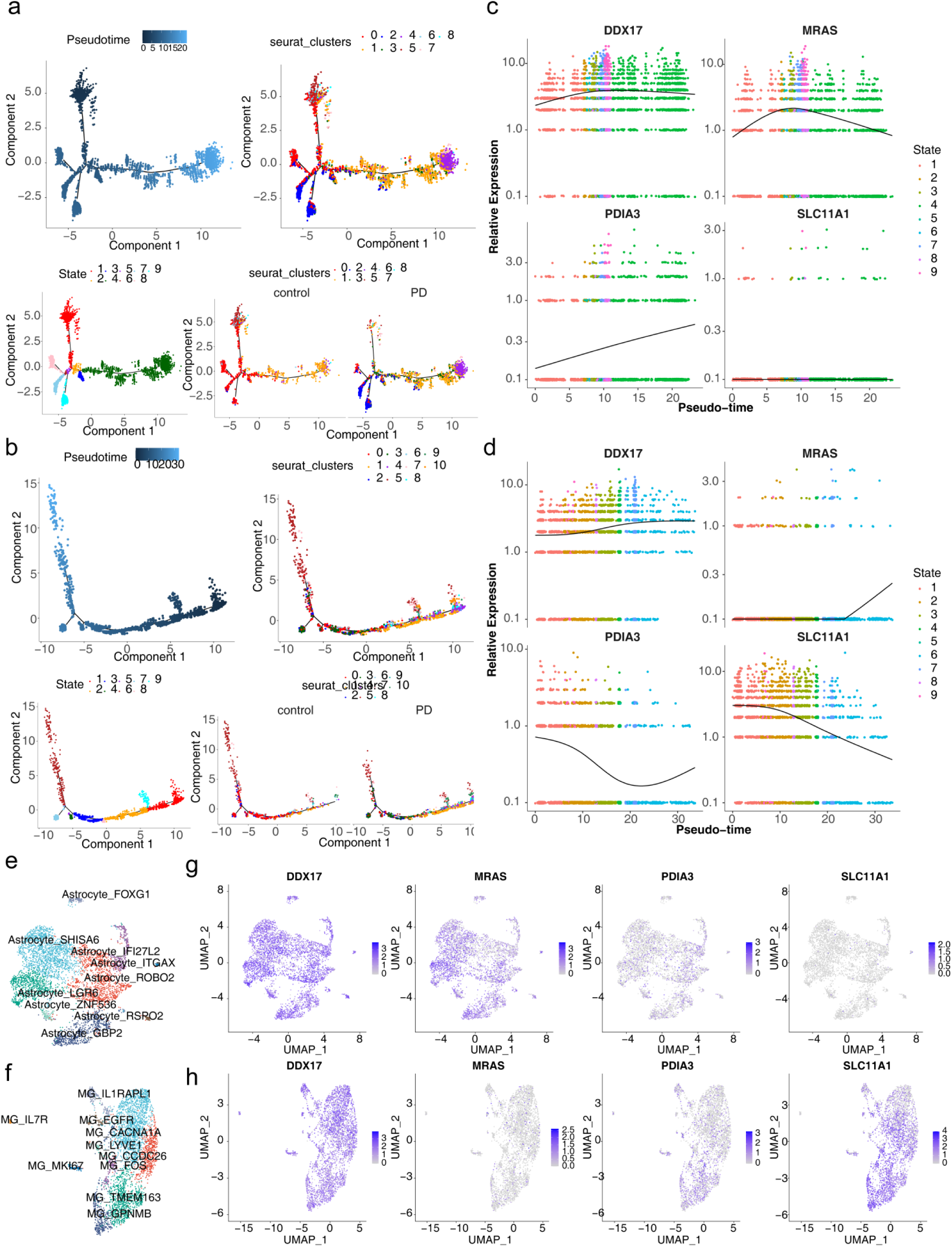
Four key genes played a role in the development of key cell sub-populations. (a,b) The expression of key genes in Astrocytes and Microglia. The UMAP plots of 9 and 11 cell clusters from Astrocytes (c) and Microglia (d). Cells from different clusters are marked by colors. The UMAP plots of 9 Astrocytes subtypes (e) and 11 Microglia subtypes (f). The UMAP plots of 9 Astrocytes subtypes (e) and 11 Microglia subtypes (f). The expression of key genes in Astrocytes subtypes (g) and Microglia subtypes (h).

The Pseudo-time series analysis tracks of Astrocytes and Microglia were divided into nine developmental stages, and there were obvious differences between PD and control groups at different developmental stages (Figures 9a and 9b). During the differentiation of astrocytes, DDX17 and MRAS first increased and then decreased, while PDIA3 continued to increase (Figure 9c). In addition, during the differentiation of microglia, DDX17 initially increased and then leveled off, MRAS increased after state 7, and PDIA3 decreased first and then increased, while SLC11a11 continued to decrease (Figure 9d).

**Figure 9.**
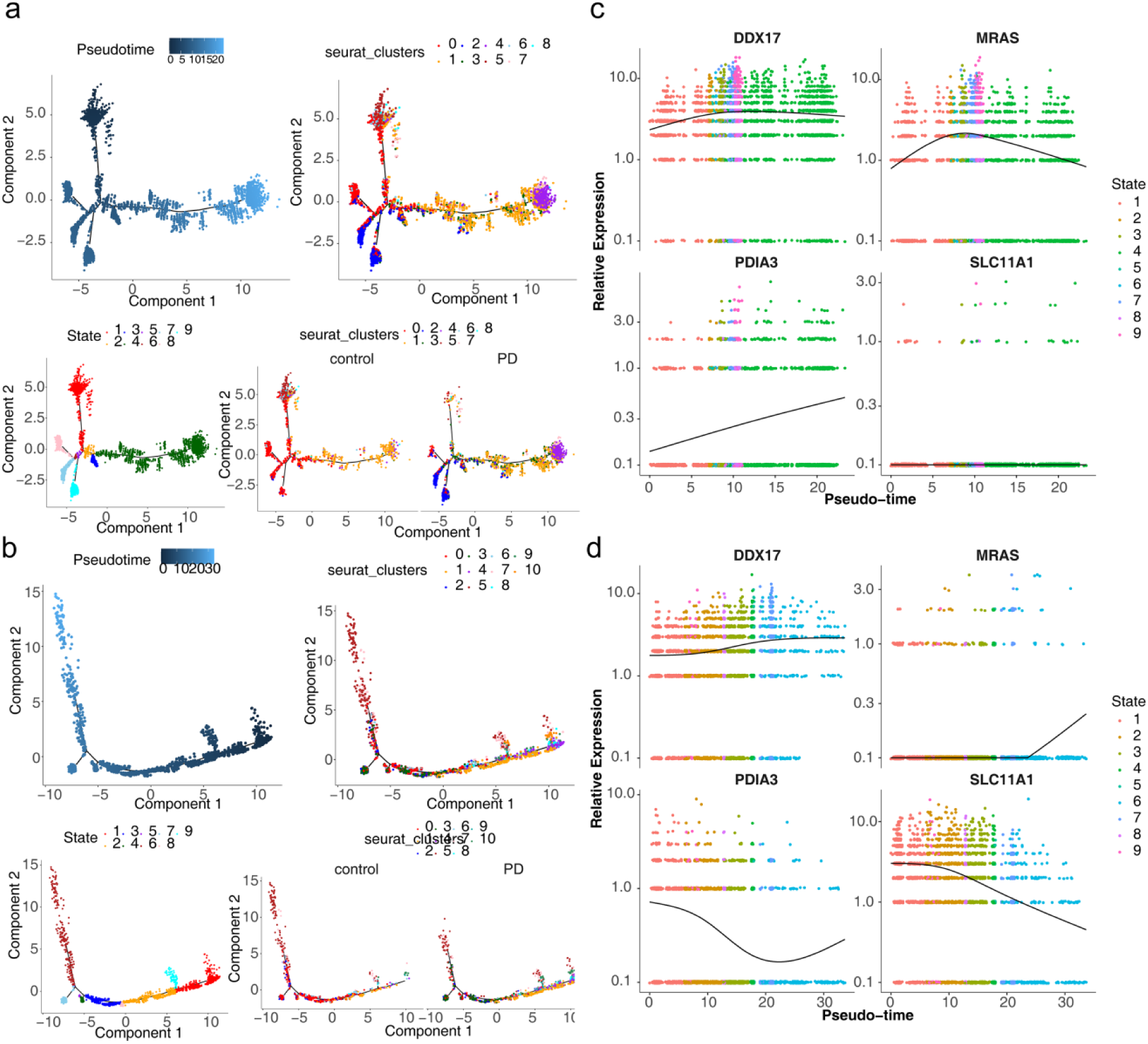
Cell pseudotiming analysis. From left to right: quasi timing trajectory map, trajectory map of different cell subsets, different differentiation trajectory stages, different differentiation trajectory stages under different groups. (a) Astrocytes and (b) Microglia. Biomarkers expression in different timing stages of (c) Astrocytes and (d) Microglia.

### Construction of key gene regulatory networks and drug prediction

The TFs most correlated with key genes in Astrocytes were BCLAF1, CHD2, and JUN, etc. In Microglia, the TFs most associated with key genes were MAX, CUX1 FLI,1, etc (Figure 10a). As shown in Figure 10b-10e, the top 10 TFs of RSS in two key cell populations and their expression in different cell populations were respectively shown. In order to further study the regulatory mechanisms involved in key genes, the lncRNA-miRNA-mRNA regulatory network was mapped. The 28 core miRNAs and 17 core lncRNAs were predicted, respectively. Finally, a regulatory network containing four key genes, 23 miRNAs, and 17 lncRNAs was constructed. For instance, XIST regulated DDX17 expression through hsa-mir-155-5p (Figure 10f). In addition, 17 drugs targeting four key genes were obtained (Figure 10g). In the drug-key genes network, ZINC CTD 00007011 was jointly forecasted by DDX17, MRAS, and PDIA3.

**Figure 10.**
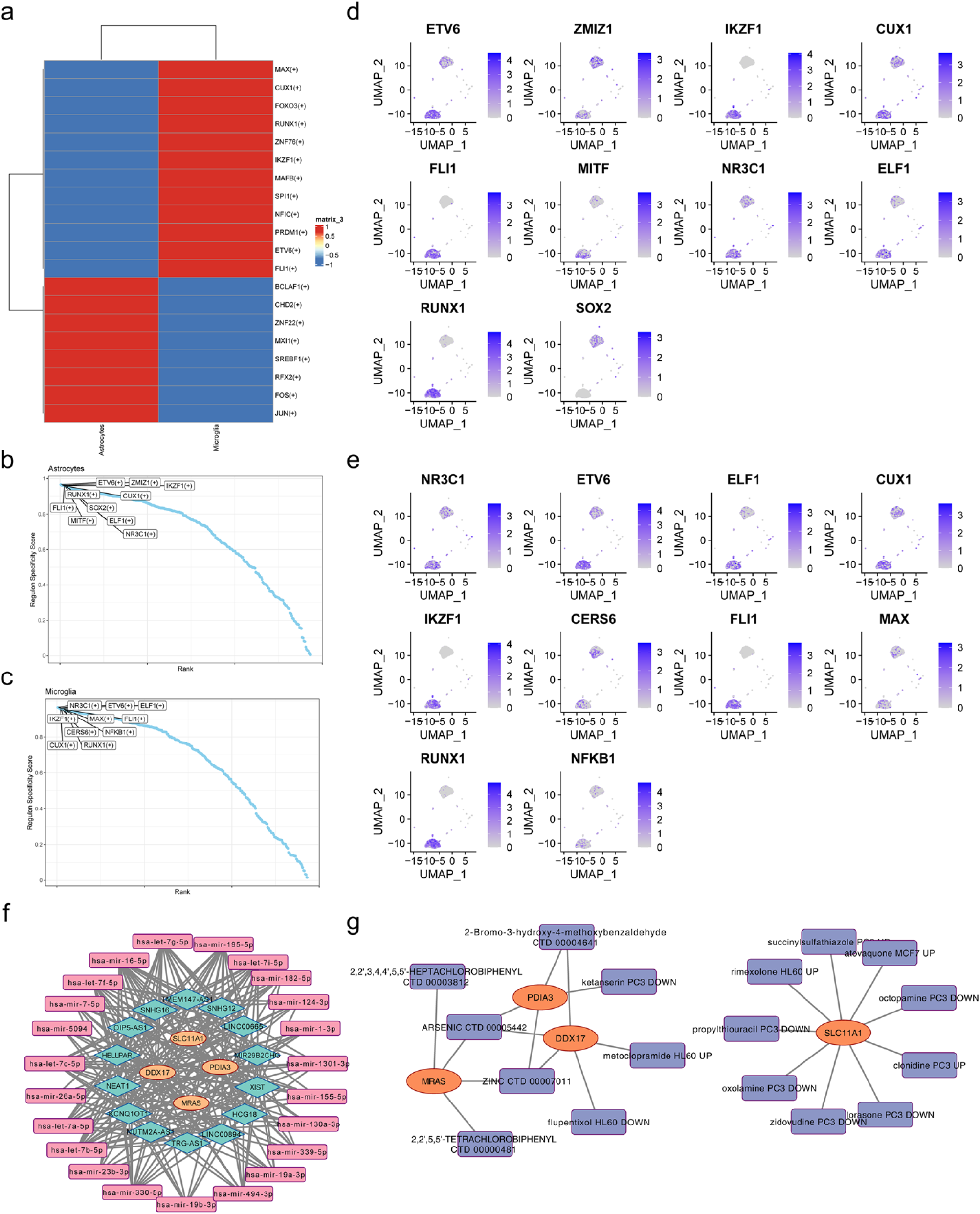
pySCENIC-analysis. (a) Prediction of transcriptional regulators of biomarkers. (b,c) Specific TF scores of the individual cell populations in Astrocytes and Microglia. (d,e) A UMAP Visualization of the Expression of Top 10 Transcription Factors (TFs) in Astrocytes and Microglia Cell Clusters. (f) lncRNA-miRNA-mRNA regulatory network of four biomarkers. (g) Drug - key genes network of four biomarkers.

### Altered expression of key genes in MPTP-induced mice

Male-specific pathogen-free (SPF) C57 mice (8-10 weeks and 25±2 g) were used in this study. The PD mouse model was induced by administering 1-methyl-4-phenyl-1,2,3,6-tetrahydropyridine (MPTP) at a dosage of 25 mg/kg through intraperitoneal injection once daily for seven consecutive days. The mice of the control group received an equivalent volume of saline solution calculated based on their body weight. Behavioral changes of the mice were monitored by rotarod test, open field test, and pole climbing performance test during the experimental period on the sixth day of MPTP injection. All animal experimental protocols were approved by the Animal Care and Use Ethics Committee of Yan’An Hospital of Kunming Medical University. (Approval NO:202301).

Compared to the control group, a significant decrease was observed in the duration of stay and the number of entries into the central zone of the open field in mice from the PD mice model group, suggesting the successful construction of the PD mice model (Figure 11a-c). Subsequently, RT-qPCR and WB validation experiments were conducted to explore the differential expression of key genes in mRNA and protein levels between PD mice and control mice. RT-qPCR results showed upregulation of DDX17 and MRAS, downregulation of SLC11a1, and no significant difference in PDIA3 expression in the PD mice model (Figure 11d). WB results demonstrated elevated expression of DDX17, MRAS, and PDIA3, as well as decreased expression of SLC11a1 in the PD mice model (Figure 11e). In addition, WB detection of DDX17-related pathway factors revealed that the expression levels of SMAD5, MYC, NFAT5, CREBBP, and DDX17 were elevated in the PD group compared with the control group (Figure 11f). These results suggested that when DDX17 expression was elevated, it might enhance the activation of SMAD5 by TGF-β signaling, and DDX17 and SMAD5 might be jointly involved in regulating the activation of glial cells and the secretion of immune factors, which further affected the inflammatory process in PD. DDX17-mediated changes in the expression of MYC might have affected the proliferation and differentiation of PD’s neural stem cells, as well as dopaminergic neuronal survival and function. Elevated DDX17 expression might have enhanced cellular responses to stresses such as osmotic stress and maintained cellular homeostasis by modulating NFAT5 activity, which in PD might have been involved in neuronal adaptive responses to oxidative stress and neurotoxins. In PD, DDX17, and CREBBP might have collectively regulated the expression of genes related to neuronal cell survival, synaptic function, etc., and had an impact on the pathological process of the disease.

**Figure 11.**
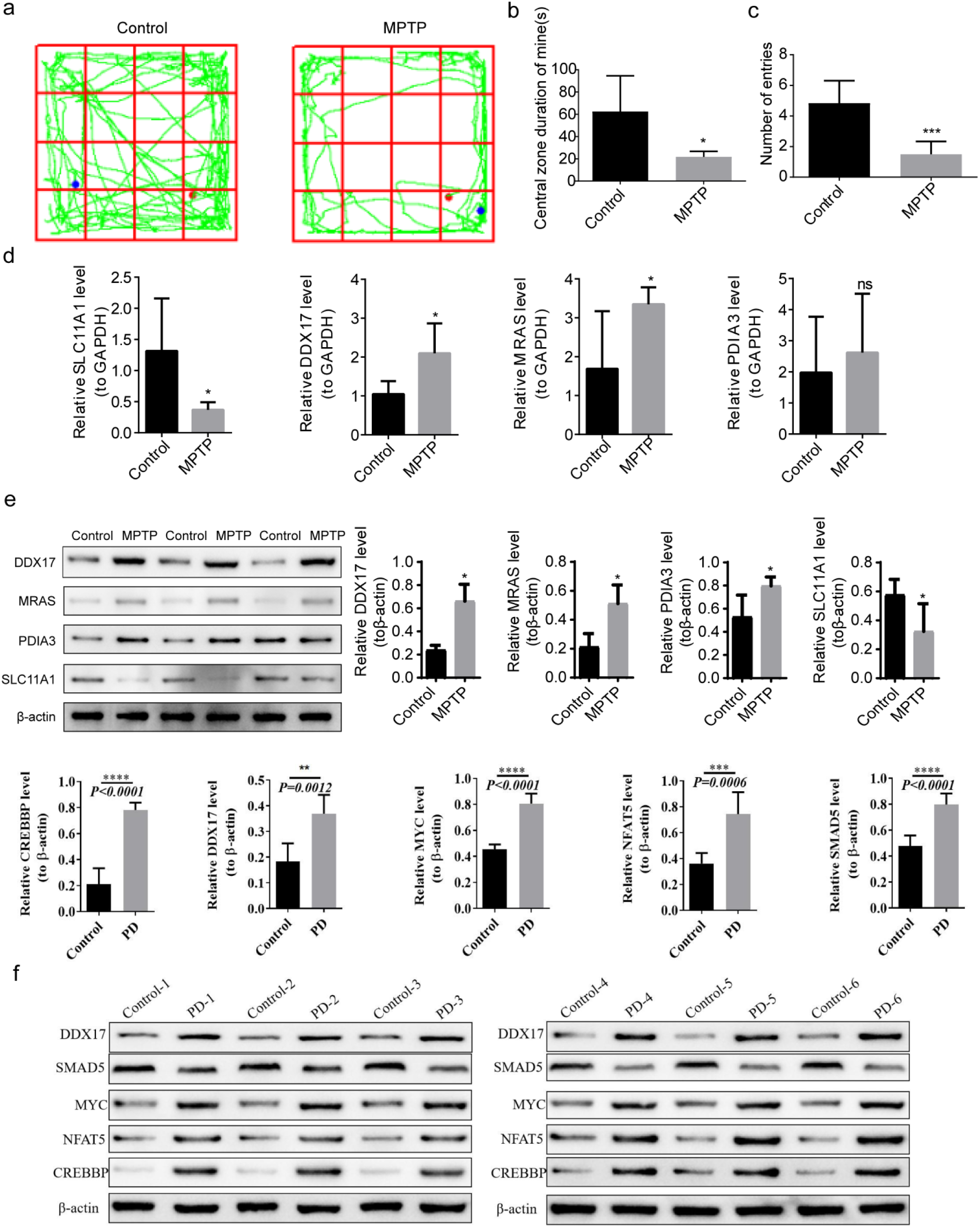
Comparison of Behavioral Phenotype and Molecular Expression in PD Mice Model Induced by MPTP. (a-c) The behavioral phenotype analysis in the open field test between the PD mice model group and the control group. (d) Comparison of RT-qPCR Expression of Key Genes in PD Mice and Normal Mice Induced by MPTP. (e) Comparison of WB Expression of Key Genes in PD Mice and Normal Mice Induced by MPTP. (f) Comparison of WB Expression of DDX17-related pathway factors in PD and controls (*\*P*< 0.05, *\*\*P*<0.01, *\*\*\*P*<0.001).

Furthermore, positive expression of GFAP, IBA1, and DDX17 was observed in the PD mice model group compared to the control group in the IF results, consistent with single-cell findings, indicating a higher proportion of Microglia and Astrocytes in PD, and higher expression of DDX17 in PD. Subsequent analysis revealed a positive co-expression of DDX17 and GFAP in the PD mice, indicating co-expression of DDX17 and Astrocytes in the PD mice model, consistent with the results of single-cell analysis (Figure 12a-f). Meanwhile, SLC11A1 showed high expression, and CD206 showed low expression in the PD group compared to the control group, which suggested that the high expression of SLC11A1 may not be directly related to the polarization of microglia during the PD process (Figure 12g). Taken together, these findings suggest that DDX17 might have influenced astrocyte-related PD development.

**Figure 12.**
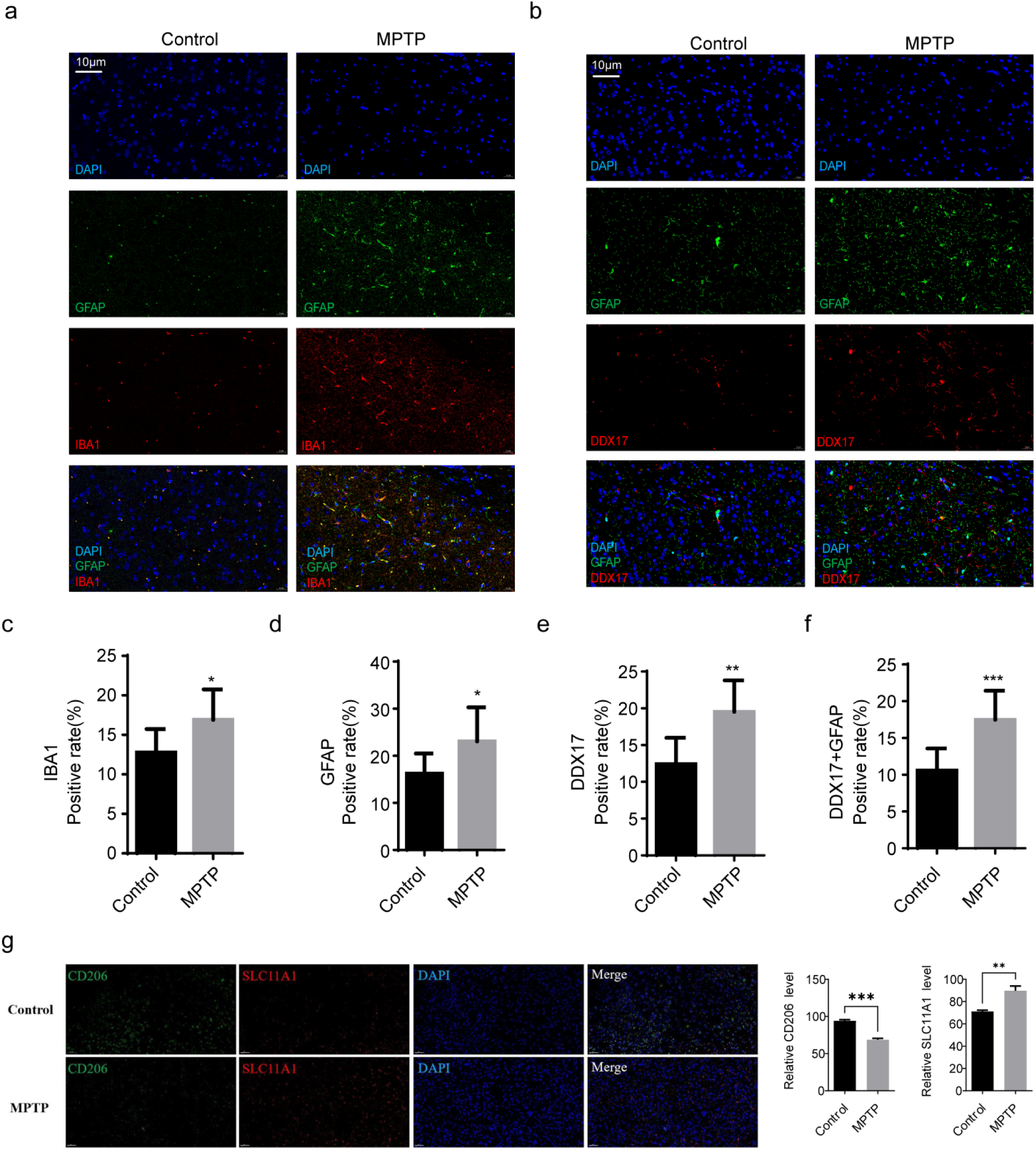
Comparison of Altered expression of key genes in PD Mice Model Induced by MPTP. (a,b) Immunofluorescence staining analysis of GFAP, IBA1, and DDX17 in PD and controls.(c-f) Gene expression positive rate analysis of GFAP, IBA1, and DDX17 and Co-expression of GFAP and DDX17 in PD Mice and Normal Mice Induced by MPTP. (g) Immunofluorescence staining analysis of SLC11A1 and CD206 in PD and controls. (*\*P*<0.05, *\*\*P*<0.01, *\*\*\*P*<0.001).

### Immunohistochemical detection

Detection of TH, α-syn expression in mouse models by immunohistochemistry. The results showed that the expressions of TH protein and α-syn protein in tissue sections were consistent, Relative to the control group, the MPTP group presented a significant downregulation of TH protein expression, while concurrently showing a remarkable upregulation of α - syn protein, and these alterations were statistically significant.(Figure 13a-b).

**Figure 13.**
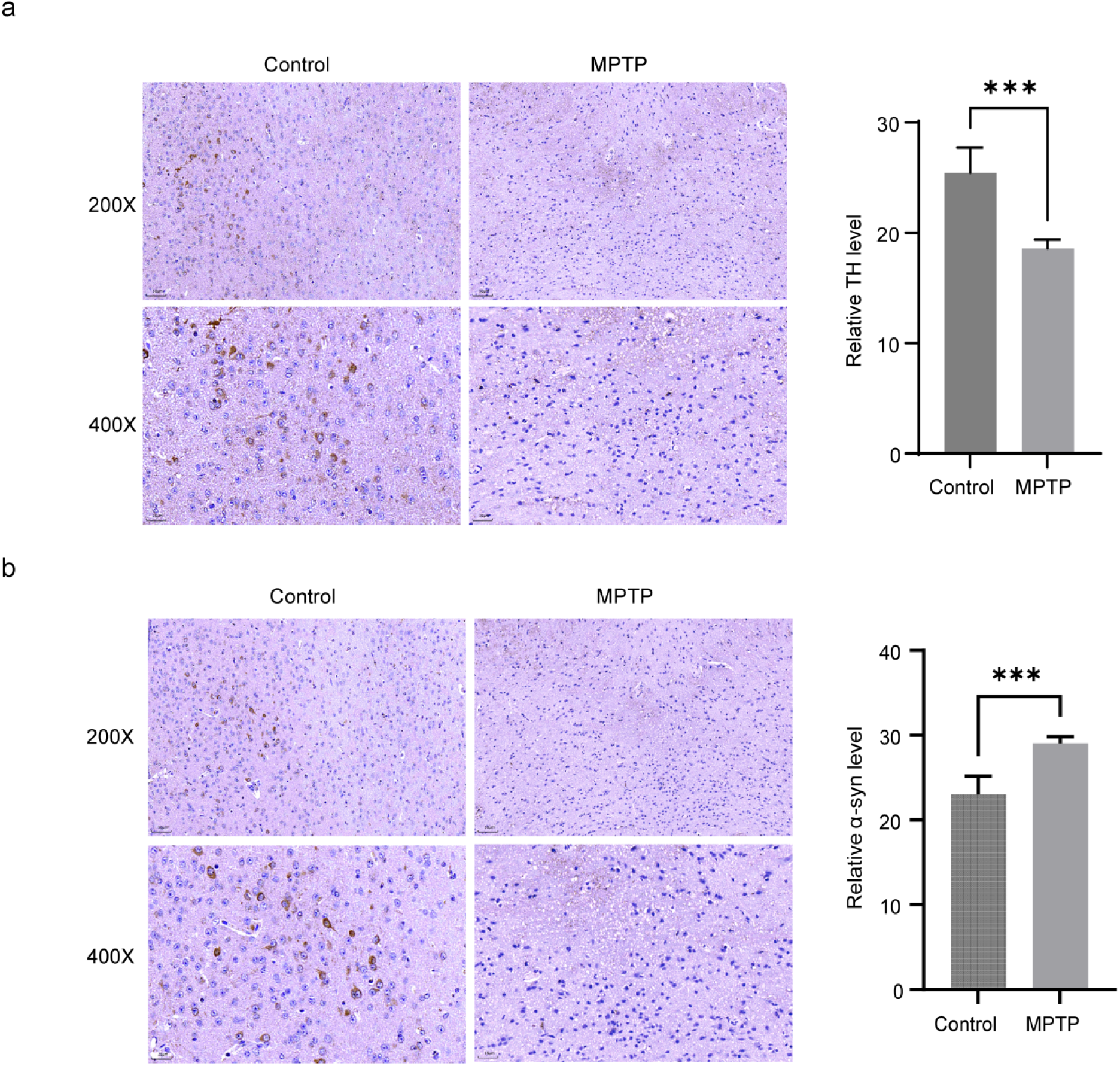
The results of immunohistochemical detection. (a) The expression of TH protein in tissue sections. (b) The expression of α-syn protein in tissue sections. *\*: P*<0.05; *\*\*\*P*<0.001.

## Discussion

PD is the second most common neurodegenerative disease in the world. With the acceleration of aging, the prevalence and incidence rate of the disease will increase gradually, but its pathogenesis has not been completely clear. More and more studies have shown that mitophagy and immunity are closely related to PD [17,18]. In this study, we identified four key genes (SLC11A1, DDX17, MRAS, and PDIA3) related to MRGs and IRGs in PD through single-cell sequencing combined with MR analysis. From a genetic perspective, we found that SLC11A1 and DDX17 are protective factors for PD, while MRAS and PDIA3 are risk factors.

SLC11A1 (solid carrier family 11 member 1) is a proton/divalent cation reverse transporter [19]. SLC11A1 encodes natural resistance associated with macrophage protein 1 (NRAMP1) and has been found to regulate macrophage activation, playing an important role in autoimmune and infectious diseases [20]. However, some studies suggest that it plays an important role in the host’s innate immunity by regulating the iron metabolism process of macrophages and affecting their early activation [21]. Li et al. [22] found a certain association between SLC11A1 gene polymorphism and tuberculosis susceptibility through Meta-Analysis, supporting the hypothesis that it may play an important role in host defense against tuberculosis development. Host genetics research has found that SLC11A1 plays an important role in neutrophil function [23]. Additionally, it has been found that the high expression of the SLC11A1 gene in gliomas is closely associated with immune activation, tumor malignancy, and the enrichment of genes related to immune therapy response [24]. Our GSEA function suggests that this gene may be involved in “antigen processing and presentation,” which further confirms its stable and important role in the immune process. In addition, we used MR for the first time to analyze the genetic role of this gene in PD and concluded that the expression of SLC11A1 is a protective factor for the occurrence of PD (OR, 0.95), that is, an increase in SLC11A1 expression may reduce the probability of PD. Therefore, SLC11A1 may play a protective role in the occurrence of PD by regulating immune processes, antigen processing, and presentation. Of course, we will continue to verify the function of this gene in PD in the future.

DDX17 (DEAD box (DDX) 17), belonging to the DEAD box RNA helicase family, is a nuclear and cytoplasmic shuttle protein that is expressed in most tissues and cells [25]. There are many studies suggesting that DDX17 is an essential factor in both cell survival and DNA repair [26]. DDX17 is considered a cofactor of miRNA microprocessors, and its dysregulation may be associated with cancer [27]. In the mechanism research of COVID-19, DDX17 has also been found to be associated with tumor cell stemness, meiosis, and DNA repair processes [28]. A new perspective suggests that DDX17 may play a role by regulating the activity of various ribonucleoprotein complexes[29]. However, there are few reports on its function in treating diseases. Our study, for the first time through MR, found that an increase in DDX17 expression may reduce the probability of PD occurrence. In addition, we found through single-cell sequencing technology that the gene has an increasing expression trend in Astrocytes and Microglia. Astrocytes and Microglia are two key regulatory factors in the inflammatory response of the central nervous system, playing important roles in some neurodegenerative diseases [30]. Sen et al. [31] believe that upregulation of calcium signaling proteins in Microglia is beneficial for effective phagocytosis of myelin sheaths, which in turn affects the nervous system. Surprisingly, we found that DDX17 is highly enriched in the ‘calcium signaling pathway.’ So, we boldly speculate that the increased expression of DDX17 in astrocytes and microglia may lead to a reduced risk of developing PD by potentially modulating the calcium signaling pathway.

MRAS (multiple RAS oncogene homolog) is a member of the RAS superfamily [32]. The GTPase of this family is considered a key regulatory factor in various cellular and developmental processes, such as differentiation, cell division, and vesicle transport [33]. MRAS is generally believed to regulate some subunits of phosphatase when combined with SHOC2 and PP1, thereby playing a role in cancer progression [34,35]. Some studies have also found that the interaction between MRAS and GNG2 may inhibit Akt and ERK activities, thereby promoting cell apoptosis and inhibiting certain cell proliferation [36]. The MR study by Saudi and Han populations found that MRAS is associated with an increased risk of coronary artery disease, obesity, and some steroid-related diseases [37]. In this study, through MR analysis, we found that the probability of PD may increase with the increase of MRAS expression, indicating that it is a risk factor for PD. In addition, we also found that the gene is overexpressed at the single-cell level in two key cell clusters associated with nerve damage: Astrocytes and Microglia. Therefore, we believe that its high expression in key cell clusters may be a factor promoting the occurrence of PD, which will provide a new approach to inhibiting negative biomarkers in the future.

PDIA3 (protein disulfide isomerase A3) is a molecular chaperone in the PDI family [38]. PDIA3 is believed to be highly expressed in cellular stress response, intercepting apoptotic cell death processes associated with endoplasmic reticulum stress and protein misfolding [39]. Our functional enrichment results showed that PDIA3 was highly enriched in ‘acute processing and presentation,’ indicating that the role of PDIA3 in the development of PD is consistent with other reports. At present, PDIA3 is increasingly being found to be involved in cancer development, and its role goes beyond its main function as a protein folding within the endoplasmic reticulum [40]. In Alzheimer’s disease (AD), PDIA3 in the cytosol interacts with other proteins to regulate the mTOR pathway, potentially leading to the upregulation of specific neurons [41]. In this study, we identified it for the first time as a potential biomarker for immune mitophagy in PD. At the single-cell level, PDIA3 shows a decreasing and then increasing expression trend in two key cell clusters, Astrocytes and Microglia. We speculate that this unstable expression may have a causal relationship with its role as a risk factor for disease. However, the specific molecular mechanisms still need to be further studied.

Single-cell data analysis showed significant up-regulation of DDX17, MRAS, and PDIA3 in astrocytes of the PD group, while SLC11A1 remained unchanged. In microglia, PDIA3 and MRAS were significantly up-regulated in the PD group, whereas SLC11A1 exhibited a down-regulation, and DDX17 did not show significant changes. Astrocytes and microglia are two major types of glial cells in the brain [42] that predominate in the central nervous system [43,44]. They play crucial roles in maintaining brain microenvironmental homeostasis, supporting neuronal function, and regulating inflammatory responses. Notably, accumulating evidence suggests that AQP4 exerts pro-inflammatory effects by promoting the release of astrocyte-derived factors that activate microglia and other astrocytes [45]. Neuroinflammation is a hallmark of PD, and astrocytes in the substantia nigra are particularly rich in AQP4. Previous studies have identified DDX17 as a regulator of AQP4 expression in astrocytes, influencing central nervous system function [46]. In our study, we observed DDX17 expression in astrocytes, suggesting its potential involvement in astrocytic regulatory mechanisms. We, therefore, hypothesize that DDX17 may modulate neuroinflammatory signaling during PD pathogenesis by influencing RNA-related functions in astrocytes. Our predicted multiple miRNAs, such as has-let-7b-5p and hsa-mir-19a-3p, have been confirmed to have significant potential roles in PD. has-let-7b-5p, under normal conditions, is involved in immune responses, cell proliferation, and differentiation. However, in PD, its sequestration by hsa_circ_0000437 may weaken its inhibitory effect on target genes, thereby exacerbating neuroinflammatory reactions [47]. On the other hand, has-miR-19a-3p, as one of the 14 specific miRNAs closely related to PD, influenced neuronal survival, apoptosis, and PD-related inflammation through intricate molecular network regulation, playing a pivotal role in the pathogenesis of PD. In terms of drugs, atovaquone has demonstrated safety in PD patients, providing preliminary evidence for its potential as a therapeutic or adjunctive treatment [48]. The significantly reduced level of octopamine in PD patients suggests its potential association with PD pathophysiology, hinting at future interventions through modulating its levels [49]. The research on propylthiouracil has revealed that its taster status is linked to increased susceptibility to PD, potentially offering a novel and simple method for early PD identification [50]. Lastly, clonidine has shown improvement in non-motor symptoms, particularly impulse control disorders, in PD patients, presenting a new option for comprehensive PD treatment [51]. These miRNAs and drugs have demonstrated diverse potential and clinical application prospects in PD research.

MPTP-induced PD mice exhibited significant motor impairments, including reduced latency time and decreased entries into the center zone in the open field test. These behavioral alterations are consistent with the motor dysfunction observed in PD patients. In this well-established mouse model, the upregulation of DDX17 and its associated pathway factors highlights its potential key role in PD pathophysiology. The SMAD5 protein is known to regulate the transforming growth factor-β (TGF-β) pathway [52], which is implicated in modulating neuroinflammation and glial cell activation [53], both critical features of PD pathogenesis. Thus, elevated SMAD5 expression in PD may influence TGF-β signaling, promoting the recruitment of microglia and astrocytes to injury sites and contributing to the chronic neuroinflammation observed in PD. Additionally, the upregulation of other DDX17-related pathway factors suggests that DDX17 may participate in the regulation of multiple signaling pathways in PD. For instance, CREBBP, a transcriptional coactivator involved in neuronal survival, synaptic function, and plasticity [54], was found to be highly expressed. This indicates that CREBBP may play a role in regulating genes essential for neuronal survival and function in PD. These findings collectively suggest that DDX17 and its associated factors may contribute to PD progression by modulating neuroinflammatory responses and neuronal viability through multiple molecular mechanisms. Subsequent validation experiments in a PD mouse model using RT-qPCR and WB confirmed the upregulation of DDX17 and MRAS, downregulation of SLC11A1, and no significant difference in PDIA3 expression at the mRNA levels. The reliability of our findings is supported by the consistency between the dataset results and experimental validation regarding DDX17 and MRAS upregulation and SLC11A1 downregulation. However, the inconsistency in PDIA3 expression between WB and RT-qPCR results may be attributed to post-transcriptional regulation and differences in protein stability.

However, this study also has some limitations. Firstly, in single-cell analysis, during the step of differential analysis, it is common for the R package Findmarker to set the built-in log_2_fc threshold at 0.25. Previous studies have also emphasized the rationality and effectiveness of this threshold in specific research contexts, further supporting its practical application [55]. Of course, we are also aware that for larger and higher-quality datasets, appropriately increasing the *|log_2_FC|* threshold may help identify more robust and significant changes in gene expression, thereby enabling more in-depth and precise exploration of biological mechanisms. In future studies, we can utilize larger and better-quality datasets while increasing the log_2_FC value to conduct more profound research. Secondly, due to limitations in research time and resources, we did not conduct direct functional experiments to validate the specific functions of these genes as protective or risk factors for PD. To more directly validate the functions of these genes, we plan to conduct a series of functional experiments in future studies. We will employ gene editing technologies (such as CRISPR/Cas9) to knock out or overexpress these genes and observe their impacts on neuronal function, inflammatory responses, and PD-related phenotypes. In addition, we can directly detect the expression of these genes in blood samples of PD patients to evaluate their feasibility as circulating biomarkers. Through these experiments, we aim to gain a deeper understanding of the roles of these genes in the pathogenesis of PD and provide new targets for PD treatment.

Based on bioinformatics methods, this study integrated scRNA seq and GWAS data to identify SLC11A1, DDX17, MRAS, and PDIA3 as potential biomarkers for neutralizing immune and mitophagy in PD. Of course, our research also has some shortcomings, such as the fact that our Mendelian randomization analysis selected a population from Europe, which may result in sample heterogeneity and lack of in vitro experimental validation or further clinical validation. We will continue to monitor the functions of these potential biomarkers in the future, hoping to provide meaningful value for the research and clinical application of PD mechanisms.

## Materials and methods

### Data Sources

The PD-related single-cell sequencing (scRNA-seq) datasets GSE157783 and GSE22491 were downloaded from the Gene Expression Omnibus (GEO) database (https://www.ncbi. nlm.nih.gov/gds). The GSE157783 dataset consisted of 5 postmortem PD midbrain tissue samples and six healthy control samples [56]. The GSE22491 dataset consisted of 10 peripheral blood mononuclear cell samples from PD patients and eight healthy control samples [57]. The 1793 IRGs were obtained from the Immunology Database and Analysis Portal (ImmPort) database (https://www.immport.org/shared/home). The 72 MRGs were collected from Kyoto Encyclopedia of Genes and Genomes (KEGG) database (https://www.kegg.jp/pathway/hsa04137) [58,59] Detailed data of exposure factor eQTL and outcome traitID for MR analysis in this study were obtained from Integrative Epidemiology Unit (IEU) Open genome-wide association study (GWAS) database (https://gwas.mrcieu.ac.uk/). A PD-related dataset (ieu-b-7) was obtained by searching for Parkinson’s disease from the IEU OpenGWAS database, including 17,891,936 single nucleotide polymorphisms (SNPs) from 482,730 samples.

### Single-cell data processing

The GSE157783 single-cell sequencing data was filtered using the CreateSeuratObject function of the Seurat package (version 4.1.0) [60] with filtering thresholds of 200< n.features <6000, n.count<20000 and percent.mt<0%. After the NormalizeData function in the Seurat package normalized the data, the FindVariableFeatures function was used to select high-variable genes based on the relationship between mean and variance [61]. After scaling the data with the ScaleData function, the statistically significant principal components (PCs) were determined by the JackStrawPlot function. Principal component analysis (PCA) was performed based on the above high variable genes and PCs. Subsequently, uniform manifold approximation and projection (UMAP) was utilized to identify different cell clusters (resolution=0.8). The cell subpopulations were annotated with reference to marker gene [62] and Single R package [63]. Cell subpopulations with significant differences between PD and control samples (p<0.05) were defined as candidate key cell subpopulations for subsequent analysis. Cell communication between candidate key cell subpopulations and other cell types was analyzed using the Cellchat R package (version 1.5.0) [64].

### Gene set variation analysis (GSVA)

In order to explore the biological pathway of differences between candidate key cell subpopulations in the GSE157783 dataset, GSVA analysis was performed on key cell subpopulations. With h.all.v2022.1.Hs.symbols.GMT GSEA Base package as background gene set, the pathway activity score assigned to individual cells. Finally, the limma package [65] was used to calculate the difference in the pathway activity score of candidate key cell subpopulations. Identification of differentially expressed genes (DEGs) The DEGs in different candidate key cell subpopulations in PD and control samples of the GSE157783 dataset were screened using the Find Marker function (|log2FC|≥0.25, *p*-value <0.05) [66]. The DEGs of two candidate key cell subpopulations were collected and combined to obtain the shared DEGs.

### Correlation analysis

To further obtain the genes with a strong correlation between IRGs and MRGs, the corrupt package was used to conduct a Spearman correlation analysis between IRGs and MRGs in two candidate key cell subpopulations [67]. The gene with |cor|>0.7 and *p*<0.01 was selected as a candidate gene for follow-up analysis. The IRGs-ARGs were derived from the intersection of candidate genes from two candidate key cell subpopulations.

### Enrichment analysis

The intersection of DEGs and IRGs-ARGs was used to obtain candidate key genes. Then, enrichment analysis was carried out to explore the biological functions and pathways enriched by the key candidate genes. The clusterProfiler package (version 4.7.1) was used for enrichment analysis based on Gene ontology (GO) and KEGG databases (*p* adj < 0.05) [68].

### MR analysis

In MR analysis, the key candidate gene was the exposure factor, and PD was the outcome. The extract_instruments function of the TwoSampleMR package was used to read exposure factors and identified SNPs that were independently related to exposure factors as instrumental variables (IVs) (*p*<5×10−8) [69]. In addition, in order to avoid the potential bias caused by the linkage disequilibrium (LD), the SNPs of the LD (r2 = 0.001, kb=100kb) were removed. Based on the GWAS data of outcome and filtered IVs, the IVs significantly correlated with the outcome were removed. The harmonise_data function of the TwoSampleMR package was utilized to unify effect alleles and effect sizes, and the exposure factors-IVs - outcomes were matched. Finally, the exposure factors that could be obtained were greater than or equal to 3 SNPs were used for analysis. In univariate MR (UVMR) analysis, Two Sample MR packages were used to carry out five algorithms to perform MR analysis (MR Egger [70], Weighted median [71], Inverse variance weighted (IVW) [72], Simple mode and Weighted mode [73]. In order to explore the causal relationship between the above exposure factors and outcome at the multivariate level, we performed multivariate MR (MVMR) analysis. The results referenced the IVW method. A sensitivity analysis was performed to test the reliability of the MR analysis. Heterogeneity was evaluated by the mr_heterogeneity function, where *p*<0.05 indicated there was heterogeneity between exposure and outcome data. The mr_pleiotropy_test function was used to evaluate horizontal pleiotropy, where *p*> 0.05 indicated no horizontal pleiotropy in MR analysis. A leave-one-out (LOO) sensitivity test was used to delete IVs one by one and determine the influence of the remaining IVs on the outcome. The positive genes of UVMR analysis were defined as key genes.

### Construction of Nomogram

Expression levels of key genes were analyzed for differences in the PD and control groups of dataset GSE22491 using the Wilcoxon rank sum test. Based on the expression levels of key genes in the training set GSE22491, with the sample’s disease status as the outcome event, a nomogram model for biomarkers was constructed using the R package “rms” (version 6.5.0). To validate the accuracy of the nomogram prediction model, a calibration curve was plotted to reflect its predictive ability. The Hosmer-Lemeshow test (HL test) was used as an indicator of model fit, with its principle being to assess the discrepancy between predicted and observed outcomes. A *p-value* greater than *0.05* indicates that the HL test is passed, suggesting no significant difference between predicted and observed values. Conversely, a p-value less than 0.05 indicates that the HL test is not passed, suggesting a significant discrepancy between predicted and observed values, thereby indicating poor model fit. To assess the predictive performance of the nomogram, the Decision Curve Analysis (DCA) curve can be plotted using the R package “media.” This analysis helps evaluate the clinical usefulness of the nomogram.

Additionally, utilizing the R package “pROC” (version 1.18.0), both the ROC curve for the nomogram and that for individual biomarkers can be plotted, with an Area Under the Curve (AUC) greater than 0.7 serving as a criterion for good predictive performance.

### Gene Set Enrichment Analysis (GSEA)

In order to explore the biological function of key genes, GSEA was performed. In the single-cell dataset, key genes were divided into high and low groups according to the median expression level, and the difference analysis was carried out. Then, all genes were sequenced according to logFC. Subsequently, c2.cp.kegg.v7.5.1.symbols. GMT was used as a background gene set, and GSEA was completed using the theclusterProfiler package (*p* adj < 0.05) [74].

### Construction of protein co-expression network

To further explore the PPI of key genes and their co-expression with other genes, a PPI network was constructed based on the Gene MANIA database (http://genemania.org/).

### Identification of key cell subpopulations and key cell subtypes

The UMAP was used to show the expression of key genes in two candidate key cell subpopulations, and the violin map was utilized to show the expression differences of key genes in two candidate key cell subpopulations of PD and control samples. Finally, the cell subpopulations with significant differences in key genes were defined as cell subpopulations. Thereafter, the cell subpopulation was annotated with reference to marker genes [75] and the SingleR package [76]. UMAP was utilized to depict the expression patterns of key genes among different key cell subtypes. Key cell subtypes with significant differences between PD and control samples (*p* <0.05) were defined as candidate key cell subtypes for subsequent analysis.

### Pseudo-time series analysis

In order to clarify the heterogeneity of key cell subpopulations in single-cell sequencing datasets, subtypes of different key cell subpopulations were re-clustered in PD and control samples, respectively, according to marker genes. Pseudo-time series analysis of different subtypes within key cell subpopulations was performed using the monocle package (version 2.22.0) to construct cell tracks [77].

### Screening of transcriptional factors (TFs)

TFs of key genes were identified from single-cell sequencing data using the pySCENIC method. In addition, TFs of key cell subpopulations were determined by regulon specificity score (RSS) [78]. UMAP visualized the expression of the top 10 RSS TFs in key cell subpopulations.

### Construction of lncRNA-miRNA-mRNA regulatory network

First, the miRNA that could interact with key genes was predicted using the miRnet database (https://www.mirnet.ca/), in which the miRNA that targeted two or more mRNAs simultaneously was used as the core miRNA. Then, lncRNAs targeting the core miRNAs were predicted by the miRnet database, and lncRNAs targeting more than 10 miRNAs at the same time were regarded as the core lncRNAs. Finally, the complete lncRNA-miRNA-mRNA regulatory network was constructed using Cytoscape software (version 3.9.1) [79].

### Drug prediction

Based on the Drug SIGnatures database (http://tanlab.ucdenver.edu/DSigDB), an enriched R package (version 3.2) was used to obtain potential drugs that key genes may target (*p* < 0.01) [80].

### Open-Field Test

Male-specific pathogen-free (SPF) C57 mice (8-10 weeks and 25±2 g) were randomly assigned to two groups. One group of mice received intraperitoneal injections of methyl-4-phenyl-1,2,3,6-tetrahydropyridine (MPTP, 25mg/kg) for six consecutive days to establish a PD mice model, while the other group received injections of physiological saline as a control. The open-field test was conducted using a manually delimited arena, which was divided into a central zone and a peripheral zone. The central zone was defined as a circular area with a radius of 25 cm centered at the midpoint of the arena, while the surrounding area constituted the peripheral zone. The mice were placed in a dark box within the arena, and after their entry into the arena, the dark box was removed, and the timing commenced. Using Smart3.0 software, the exploratory trajectories, distances traveled, speeds, and the number of entries into the central zone during a 10-minute period were observed and recorded [81,82]. Throughout the course of the experiments, no symptoms indicative of severe illness or mortality were observed in the animals.

### Real-time quantitative polymerase chain reaction (RT-qPCR)

Brain tissue samples from the substantia nigra region were collected at 50 mg each from Parkinson’s disease mice and control mice, and RT-qPCR analysis was performed to compare the differences in mRNA expression between the disease and control groups. Total RNA from the samples was isolated and purified using the TRIzol method according to the manufacturer’s protocol. The RNA was then reverse-transcribed into cDNA using the SureScript-First-strand-cDNA-synthesis-kit. The cDNA was diluted 5-fold and mixed with 5×BlazeTaq qPCR Mix and primers (**Table 1**) for target genes. The RT-qPCR amplification was performed with the following program: initial denaturation at 95°C for 1 min, denaturation at 95°C for 20 s, annealing at 55°C for 20 s, and extension at 72°C for 30 s for 40 cycles. The data analysis was conducted using the 2^−ΔΔCt^ method, and the results were normalized to GAPDH as an internal control in accordance [83].

**Table 1.**
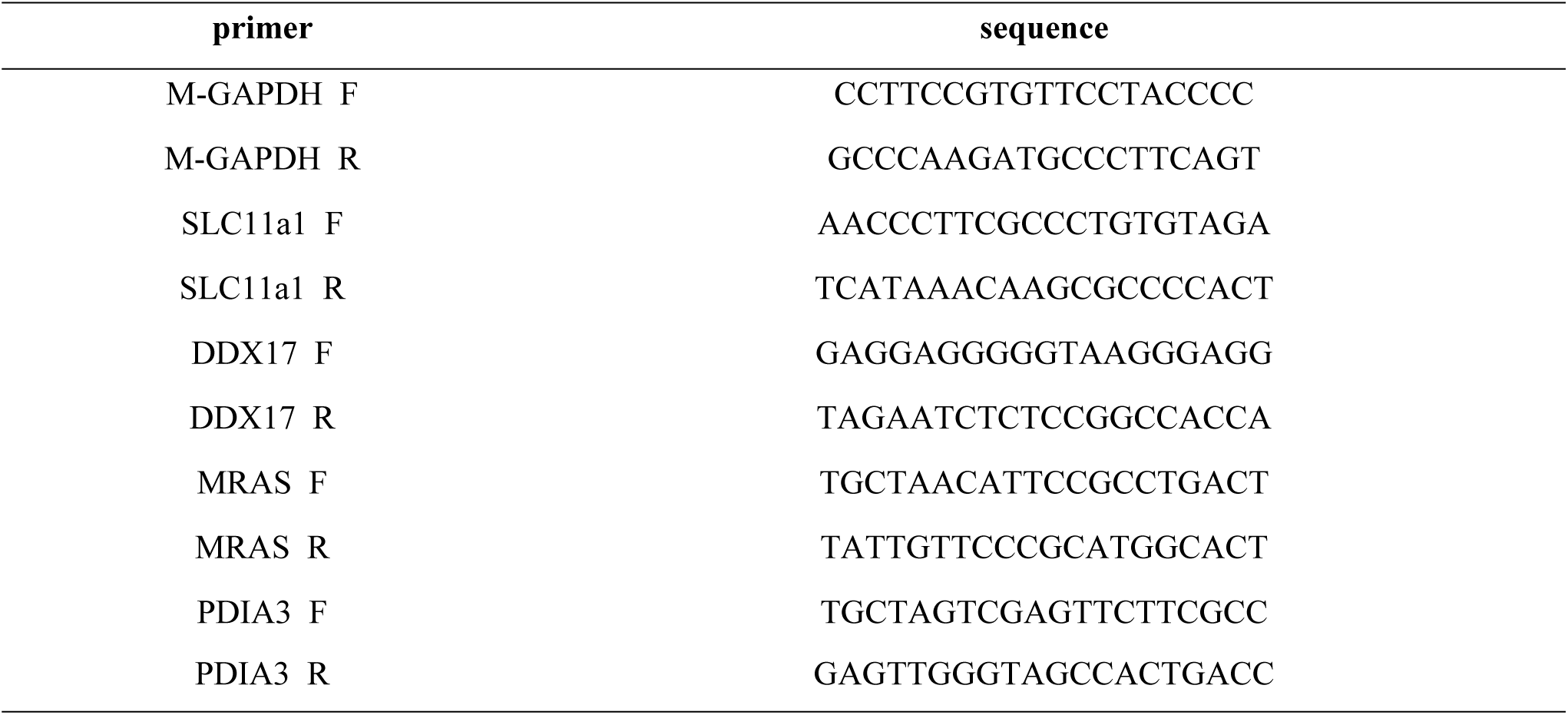
The list of qRT-PCR primers.

### Western blot (WB)

RIPA lysis buffer (Servicebio) containing a protease inhibitor (Servicebio) was used to lyse 100 mg of brain substantia nigra tissue, and the protein concentration was measured using the BCA protein assay kit (Beyotime). Subsequently, the proteins were separated by 10% SDS-PAGE and transferred to a PVDF membrane (Millipore). The membrane was blocked with 5% BSA (Solarbio) and then incubated with the corresponding primary antibodies. Subsequent incubation involved secondary antibodies, followed by color development using a highly sensitive ECL chemiluminescence kit (Affinity). The imaging was performed using the Vilber Lourmat chemiluminescence imaging system, and images were captured. Band intensity was quantified using image analysis software (ImageJ) [84]. The antibodies used in this study are shown in **Table 2**.

**Table 2.** The antibodies used in this study.

| name | manufacturer | No. | dilution |
| --- | --- | --- | --- |
| SLC11a1 | abcam | ab211448 | 1:1000 |
| DDX17 | abcam | ab180190 | 1:1000 |
| MRAS | abcam | ab1765 70 | 1:1000 |
| PDIA3 | Affinity | DF6230 | 1:2000 |
| β-Actin | Proteintech | 66009-1-Ig | 1:25000 |
| Goat anti-rabbit IgG labeled with HRP | invitrogen | AB 228338 | 1:2500 |
| Goat anti-mouse IgG labeled with HRP | Servicebio | GB23301 | 1:5000 |

**Table 3.**
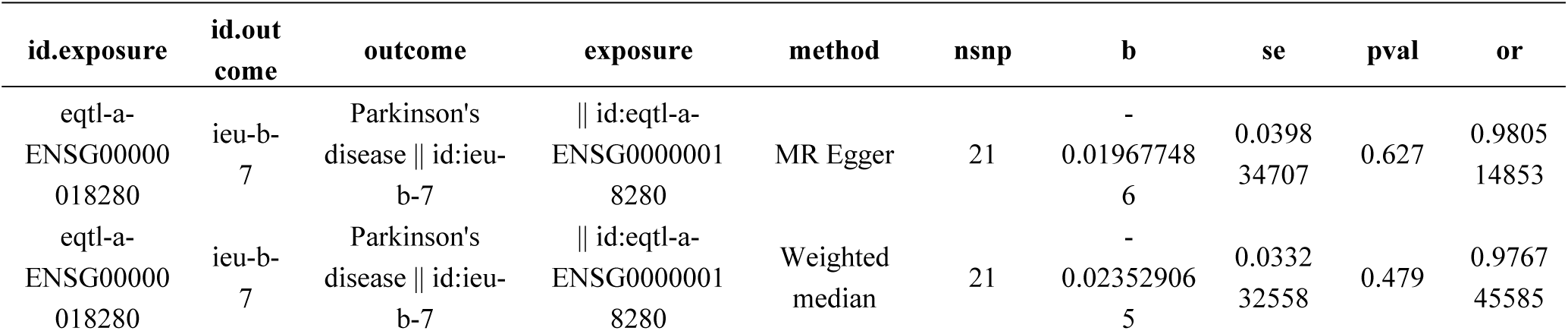
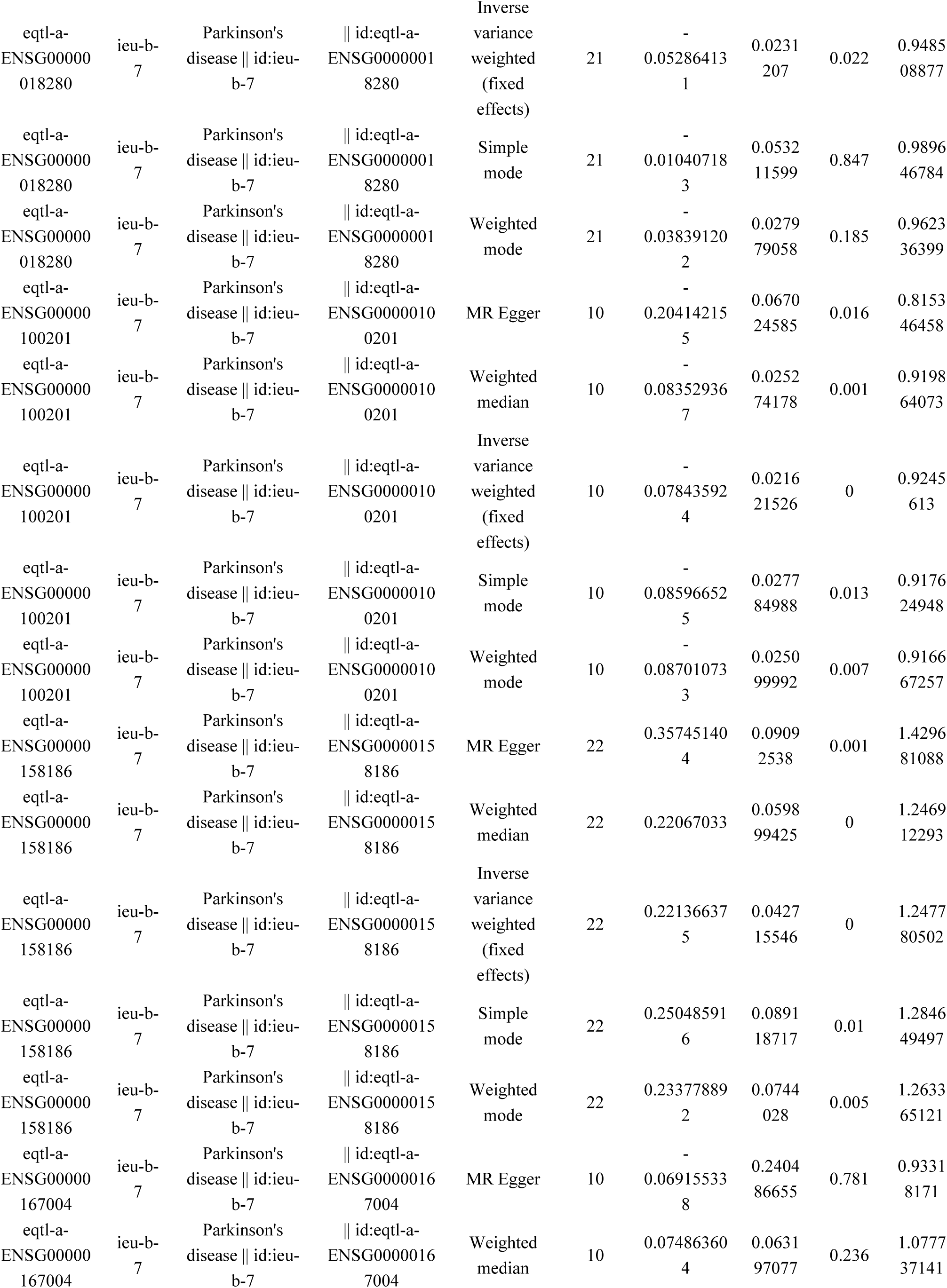
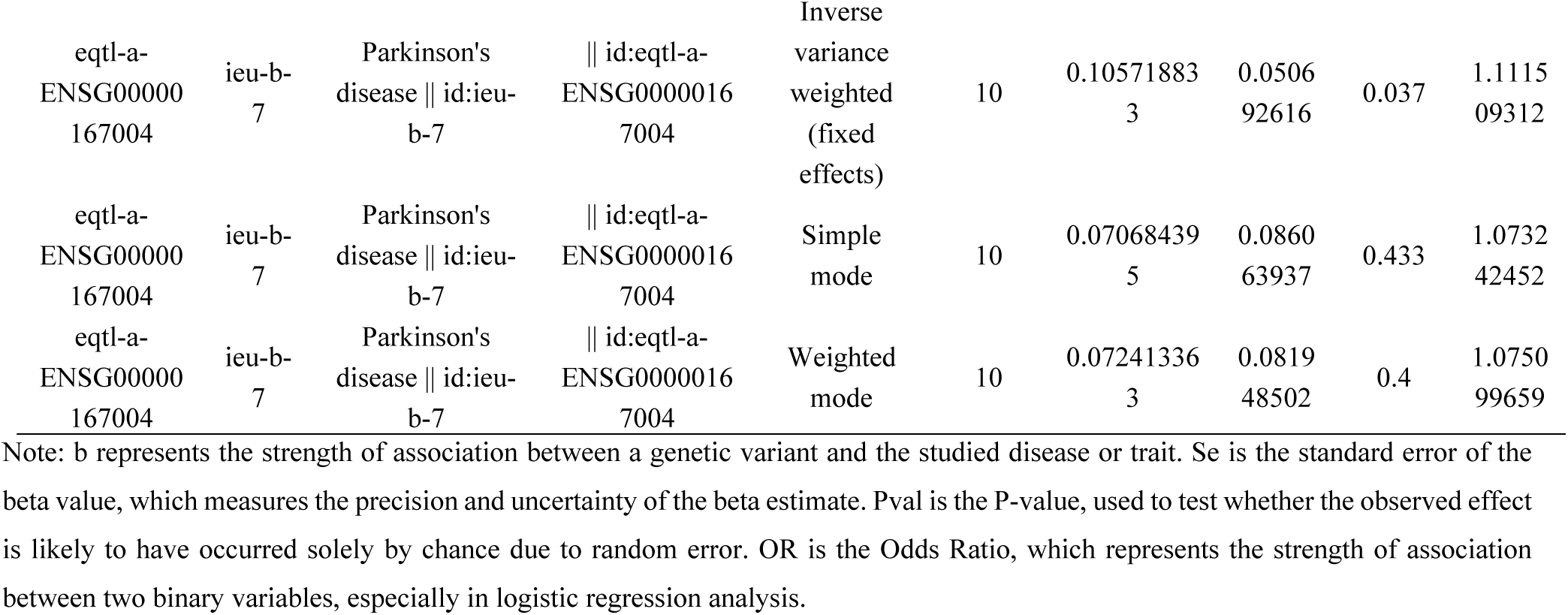
UVMR analysis of SLC11a1, DDX17, MRAS and PDIA3 and PD.

**Table 4.**
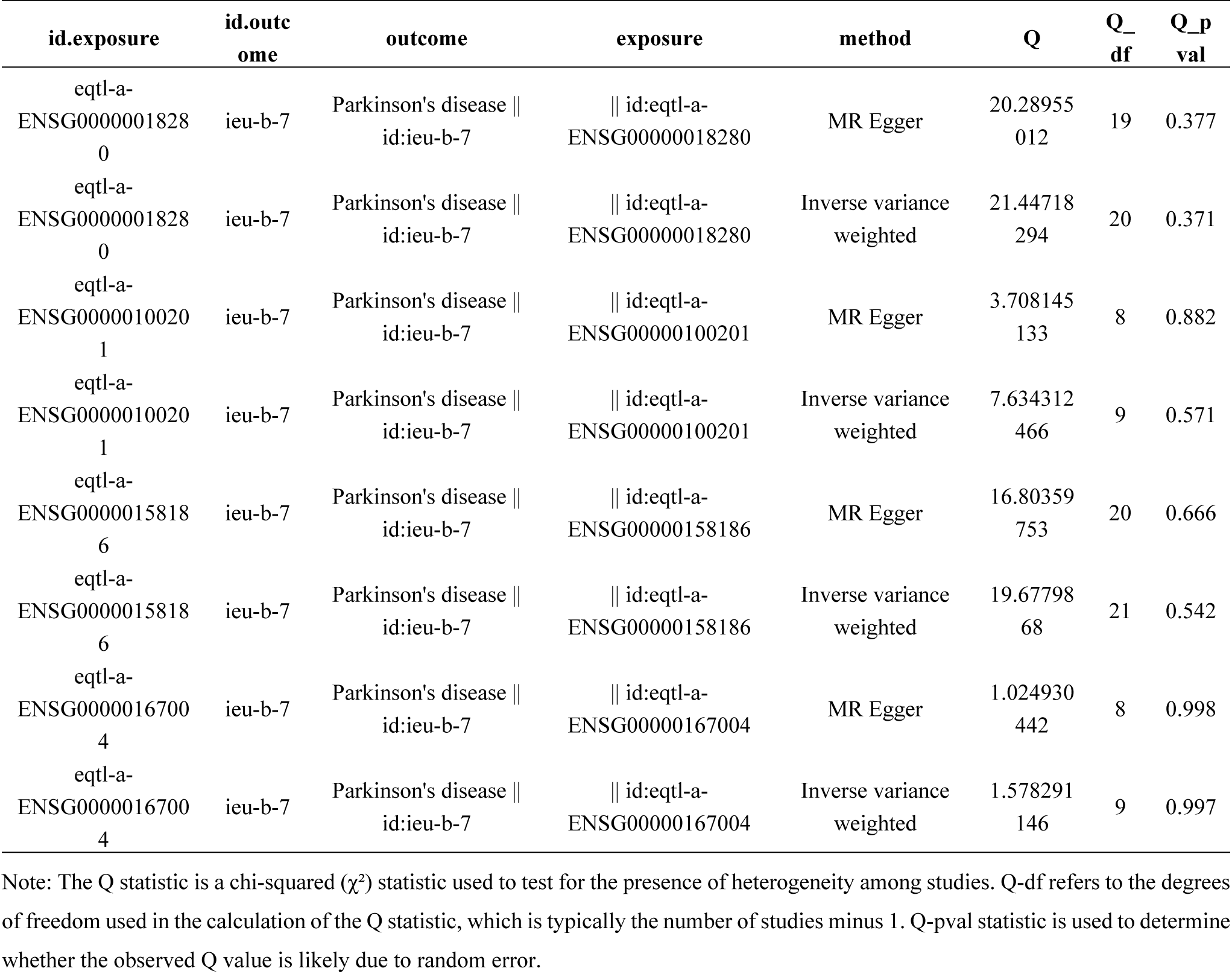
Heterogeneity analysis between exposure and outcome datasets.

| id.exposure | id.outc<br>ome | outcome | exposure | method | Q | Q_<br>df | Q_p<br>val |
| --- | --- | --- | --- | --- | --- | --- | --- |
| eqtl-a-ENSG0000001828<br>0 | ieu-b-7 | Parkinson's disease \|\|<br>id:ieu-b-7 | \|\| id:eqtl-a-ENSG00000018280 | MR Egger | 20.28955<br>012 | 19 | 0.377 |
| eqtl-a-ENSG0000001828<br>0 | ieu-b-7 | Parkinson's disease \|\|<br>id:ieu-b-7 | \|\| id:eqtl-a-ENSG00000018280 | Inverse variance<br>weighted | 21.44718<br>294 | 20 | 0.371 |
| eqtl-a-ENSG0000010020<br>1 | ieu-b-7 | Parkinson's disease \|\|<br>id:ieu-b-7 | \|\| id:eqtl-a-ENSG00000100201 | MR Egger | 3.708145<br>133 | 8 | 0.882 |
| eqtl-a-ENSG0000010020<br>1 | ieu-b-7 | Parkinson's disease \|\|<br>id:ieu-b-7 | \|\| id:eqtl-a-ENSG00000100201 | Inverse variance<br>weighted | 7.634312<br>466 | 9 | 0.571 |
| eqtl-a-ENSG0000015818<br>6 | ieu-b-7 | Parkinson's disease \|\|<br>id:ieu-b-7 | \|\| id:eqtl-a-ENSG00000158186 | MR Egger | 16.80359<br>753 | 20 | 0.666 |
| eqtl-a-ENSG0000015818<br>6 | ieu-b-7 | Parkinson's disease \|\|<br>id:ieu-b-7 | \|\| id:eqtl-a-ENSG00000158186 | Inverse variance<br>weighted | 19.67798<br>68 | 21 | 0.542 |
| eqtl-a-ENSG0000016700<br>4 | ieu-b-7 | Parkinson's disease \|\|<br>id:ieu-b-7 | \|\| id:eqtl-a-ENSG00000167004 | MR Egger | 1.024930<br>442 | 8 | 0.998 |
| eqtl-a-ENSG0000016700<br>4 | ieu-b-7 | Parkinson's disease \|\|<br>id:ieu-b-7 | \|\| id:eqtl-a-ENSG00000167004 | Inverse variance<br>weighted | 1.578291<br>146 | 9 | 0.997 |

Note: The Q statistic is a chi-squared ( $\chi^2$ ) statistic used to test for the presence of heterogeneity among studies. Q-df refers to the degrees of freedom used in the calculation of the Q statistic, which is typically the number of studies minus 1. Q-pval statistic is used to determine whether the observed Q value is likely due to random error.

**Table 5.** Horizontal pleiotropy analysis between exposure and outcome datasets.

| id.exposure | id.outco<br>me | outcome | exposure | egger_inter<br>cept | se | pval | jud<br>ge |
| --- | --- | --- | --- | --- | --- | --- | --- |
| eqtl-a-ENSG0000001828<br>0 | ieu-b-7 | Parkinson's disease \|\|<br>id:ieu-b-7 | \|\| id:eqtl-a-ENSG00000018280 | -<br>0.01025084<br>1 | 0.00984<br>5403 | 0.31086<br>0109 | NO |
| eqtl-a-ENSG00000010020<br>1 | ieu-b-7 | Parkinson's disease \|\|<br>id:ieu-b-7 | \|\| id:eqtl-a-ENSG000000100201 | 0.04073765<br>9 | 0.02055<br>9458 | 0.08286<br>18 | NO |
| eqtl-a-ENSG00000015818<br>6 | ieu-b-7 | Parkinson's disease \|\|<br>id:ieu-b-7 | \|\| id:eqtl-a-ENSG000000158186 | -<br>0.03022883<br>2 | 0.01782<br>9886 | 0.10551<br>2499 | NO |
| eqtl-a-ENSG00000016700<br>4 | ieu-b-7 | Parkinson's disease \|\|<br>id:ieu-b-7 | \|\| id:eqtl-a-ENSG000000167004 | 0.02274997<br>3 | 0.03058<br>2763 | 0.47822<br>4574 | NO |

Note: SE stands for the standard error of an estimated parameter. Pval represents the significance level in hypothesis testing.

### Immunofluorescence (IF) staining

Appropriate amounts of brain substantia nigra tissues from PD mice and control mice were fixed overnight in 4% paraformaldehyde, dehydrated in an ethanol gradient, and embedded in paraffin. The tissues were sectioned at a thickness of 5 μm using a microtome, baked at 64°C for 1 hour, deparaffinized in xylene, and subjected to antigen retrieval using high pressure after hydration in an ethanol gradient. Subsequently, the tissues were incubated with 5% bovine serum albumin at 37°C for 30 min, followed by primary antibody GFAP (Affinity, 1:50) incubation overnight at 4°C, and a 30-min re-warming step at 37°C the next day. After washing with PBS three times for 5 min each, the tissues were treated with the corresponding secondary antibody at 37°C for 30 min, washed with PBS three times for 5 min each, and then incubated with primary antibodies IBA1 (Affinity, 1:50) and DDX17 (Proteintech, 1:20), LC11A1 (Affinity, DF12509, 1:100), and CD206 (Proteintech, 18704-1-AP, 1:100). DAPI staining was performed by adding DAPI solution to the cells, incubating at room temperature in the dark for 5 min, and washing with PBS three times for 5 min each. Subsequently, images were captured using a light microscope. Five fields of view were selected for imaging, and the positivity rate was calculated [85].

### Immunohistochemical

Appropriate amounts of brain substantia nigra tissues from PD mice and control mice were fixed overnight in 4% paraformaldehyde, dehydrated in an ethanol gradient. Following dehydration, the tissue underwent clearing in xylene to facilitate paraffin infiltration. Subsequently, the tissue block was immersed in paraffin multiple times to ensure thorough impregnation. Once the paraffin infiltration was complete, the tissue block was embedded in a mold to form a paraffin block. During slicing, the paraffin block was cut into 3μm-thick sections using a microtome, which was then flattened through alcohol and a water bath before being attached to glass slides. The slides were then baked at 64°C to fix the sections in place. Next, the primary antibody incubation process began with slide preparation steps, which included baking, dewaxing, and hydration. Antigen retrieval was then performed to expose the antigenic epitopes. After blocking endogenous peroxidase activity with 3% H_2_O_2_, non-specific binding sites were sealed with 5% bovine serum albumin (BSA) (Solarbio, Beijing, China). The primary antibody was diluted with 2% BSA according to the manufacturer’s instructions and applied to the slides, which were then incubated overnight at 4°C. On the following day, secondary antibody incubation was carried out, involving rewarming the slides, adding a reaction enhancer solution, and applying an enhanced enzyme-labeled goat anti-mouse/rabbit IgG polymer. DAB staining and hematoxylin counterstaining were then performed to visualize the target protein and cell nuclei, respectively. After staining, the slides underwent dehydration and clearing before being mounted with neutral gum and subjected to whole-slide scanning. The results were analyzed using ImageJ-pro-plus software, and bar graphs were created for statistical purposes using GraphPad Prism.

### Statistical analysis

Statistical analysis was performed using Prism GraphPad, with the experimental data presented as mean ± standard error of the mean (*Mean ± SEM)*. Student’s t-test and Wilcoxon rank-sum test were used for intergroup comparisons, while one-way analysis of variance (ANOVA) was employed for comparisons involving multiple groups. The R software was used to process and analyze the single cell and transcriptome data. The p-value < 0.05 was considered statistically significant.

### Ethics approval statement

This study was performed in line with the principles of the Declaration of Helsinki. Approval was granted by the Ethics Committee of Yan’An Hospital of Kunming Medical University. (Approval NO:202301).

## Acknowledgements

We would like to express our sincere gratitude to all individuals and organizations who supported and assisted us throughout this research. Special thanks to: Research Fund of Yunnan Provincial Department of Education (2024J0284) and Kunming Health Technology Talent Fund (2022-SW-07). In conclusion, we extend our thanks to everyone who has supported and assisted us along the way. Without their support, this research would not have been possible.

## Author Contributions

**Conceptualization**: Xiaolin Dong

**Data curation**: Gang Wu, Lijuan Zhang, Yifan Ling

**formal analysis**: Lijuan Zhang, Qingyun Li

**Funding acquisition**: Xiaolin Dong

**Investigation**: Xiaolin Dong, Qingyun Li, Yifan Ling

**Methodology**: Xiaolin Dong, Yanping Li, Furong Jin

**Project administration**: Yanming Xu

**Software**: Gang Wu, Furong Jin

**Supervision**: Yanming Xu:

**Visualization:** Yanping Li, Rui Li, Yifan Ling

**Writing – original draft**: Xiaolin Dong

**Writing – review & editing**: Xiaolin Dong, Gang Wu, Yanming Xu:

All authors agree to be accountable for the content of the work.

## Data Availability Statement

The raw data could be obtained on request from the corresponding author. The datasets analysed during the current study were available in the [GEO] repository, [https://www.ncbi.nlm.nih.gov/geo/]; [ImmPort] repository, [https://www.immport.org/shared/home] and [IEU OpenGWAS] repository, [https://gwas.mrcieu.ac.uk/].

## Supporting information

**s1 Fig Single cell annotation**. (**A**) From left to right, the distribution of nFeature_RNA (the number of genes measured in each cell), nCount_RNA (the sum of all gene expression measured in each cell), and percent.mt (the proportion of mitochondrial genes measured in each cell) in each sample before quality control is shown. (**B**) From left to right, the distribution of nFeature_RNA, nCount_RNA, and percent.mt after quality control in each sample after quality control is shown.

**S2 Fig Cell communication**. (A) Detailed network for each cell interaction. (B) Ligand-receptor interaction for cellular communication

**S1 Table** Receptor-ligand interactions in candidate key cell subpopulations.

**S2-S3 Table** the signaling pathways for differences between two candidate key cell subpopulations

**S4 Table** MR analysis estimates for the association between candidate key genes and PD.

## References

[1] Winblad B, Amouyel P, Andrieu S, Ballard C, Brayne C, Brodaty H, et al. Defeating Alzheimer’s disease and other dementias: a priority for European science and society. Lancet Neurol. 2016;15(5):455–532.

[2] Chandra R, Hiniker A, Kuo YM, Nussbaum RL, Liddle RA. α-Synuclein in gut endocrine cells and its implications for Parkinson’s disease. JCI Insight. 2017;2(12):e92295.

[3] Kalia LV, Lang AE. Parkinson disease in 2015: Evolving basic, pathological and clinical concepts in PD. Nat Rev Neurol. 2016;12(2):65-6.

[4] Dorsey ER, Sherer T, Okun MS, Bloem BR. The Emerging Evidence of the Parkinson Pandemic. J Parkinsons Dis. 2018;8(s1):S3–8.

[5] Zhou Y, Li Z, Chi C, Li C, Yang M, Liu B. Identification of Hub Genes and Potential Molecular Pathogenesis in Substantia Nigra in Parkinson’s Disease via Bioinformatics Analysis. Parkinsons Dis. 2023;2023:6755569.

[6] Liu X, Chen J, Guan T, Yao H, Zhang W, Guan Z, et al. Correction to: miRNAs and target genes in the blood as biomarkers for the early diagnosis of Parkinson’s disease. BMC Syst Biol. 2019;13(1):20.

[7] Waldner A, Dassati S, Redl B, Smania N, Gandolfi M. Apolipoprotein D Concentration in Human Plasma during Aging and in Parkinson’s Disease: A Cross-Sectional Study. Parkinsons Dis. 2018;2018:3751516.

[8] Horsager J, Borghammer P. Brain-first vs. body-first Parkinson’s disease: An update on recent evidence. Parkinsonism Relat Disord. 2024;122:106101.

[9] Kamath T, Abdulraouf A, Burris SJ, Langlieb J, Gazestani V, Nadaf NM, et al. Single-cell genomic profiling of human dopamine neurons identifies a population that selectively degenerates in Parkinson’s disease. Nat Neurosci. 2022;25(5):588–95.

[10] Pissadaki EK, Bolam JP. The energy cost of action potential propagation in dopamine neurons: clues to susceptibility in Parkinson’s disease. Front Comput Neurosci. 2013;7:13.

[11] Rani L, Sahu MR, Mondal AC. Age-related Mitochondrial Dysfunction in Parkinson’s Disease: New Insights Into the Disease Pathology. Neuroscience. 2022;499:152–69.

[12] Yang Z, Goronzy JJ, Weyand CM. Autophagy in autoimmune disease. J Mol Med (Berl). 2015;93(7):707–717.

[13] Wang L, Das JK, Kumar A, Peng HY, Ren Y, Xiong X, et al. Autophagy in T-cell differentiation, survival and memory. Immunol Cell Biol. 2021;99(4):351–60.

[14] Li X, Sundquist J, Sundquist K. Subsequent risks of Parkinson disease in patients with autoimmune and related disorders: a nationwide epidemiological study from Sweden. Neurodegener Dis. 2012;10(1-4):277–84.

[15] Patani R, Hardingham GE, Liddelow SA. Functional roles of reactive astrocytes in neuroinflammation and neurodegeneration. Nat Rev Neurol. 2023;19(7):395–409.

[16] Xin G, Niu J, Tian Q, Fu Y, Chen L, Yi T, et al. Identification of potential immune-related hub genes in Parkinson’s disease based on machine learning and development and validation of a diagnostic classification model. PLoS One. 2023;18(12):e0294984.

[17] Dernie F. Mitophagy in Parkinson’s disease: From pathogenesis to treatment target. Neurochem Int. 2020;138:104756.

[18] Wang W, Wang X, Fujioka H, Hoppel C, Whone AL, Caldwell MA, et al. Parkinson’s disease-associated mutant VPS35 causes mitochondrial dysfunction by recycling DLP1 complexes. Nat Med. 2016;22(1):54–63.

[19] Blackwell JM, Searle S, Mohamed H, White JK. Divalent cation transport and susceptibility to infectious and autoimmune disease: continuation of the Ity/Lsh/Bcg/Nramp1/Slc11a1 gene story. Immunol Lett. 2003;85(2):197–203.

[20] Blackwell JM, Goswami T, Evans CA, Sibthorpe D, Papo N, White JK, et al. SLC11A1 (formerly NRAMP1) and disease resistance. Cell Microbiol. 2001;3(12):773–84.

[21] Wyllie S, Seu P, Goss JA. The natural resistance-associated macrophage protein 1 Slc11a1 (formerly Nramp1) and iron metabolism in macrophages. Microbes Infect. 2002;4(3):351–359. doi:10.1016/s1286-4579(02)01548-4

[22] Li X, Yang Y, Zhou F, Zhang Y, Lu H, Jin Q, et al. SLC11A1 (NRAMP1) polymorphisms and tuberculosis susceptibility: updated systematic review and meta-analysis. PLoS One. 2011;6(1):e15831.

[23] Braliou GG, Kontou PI, Boleti H, Bagos PG. Susceptibility to leishmaniasis is affected by host SLC11A1 gene polymorphisms: a systematic review and meta-analysis. Parasitol Res. 2019;118(8):2329–42.

[24] Xu H, Zhang A, Fang C, Zhu Q, Wang W, Liu Y, et al. SLC11A1 as a stratification indicator for immunotherapy or chemotherapy in patients with glioma. Front Immunol. 2022;13:980378.

[25] Xu K, Sun S, Yan M, Cui J, Yang Y, Li W, et al. Corrigendum: DDX5 and DDX17-multifaceted proteins in the regulation of tumorigenesis and tumor progression. Front Oncol. 2023;13:1204712.

[26] Bader AS, Luessing J, Hawley BR, Skalka GL, Lu WT, Lowndes NF, et al. DDX17 is required for efficient DSB repair at DNA:RNA hybrid deficient loci. Nucleic Acids Res. 2022;50(18):10487–502.

[27] Wu KJ. The role of miRNA biogenesis and DDX17 in tumorigenesis and cancer stemness. Biomed J. 2020;43(2):107–14.

[28] Häfner SJ. Bargain with the tooth fairy - The savings accounts for dental stem cells. Biomed J. 2020;43(2):99–106.

[29] Giraud G, Terrone S, Bourgeois CF. Functions of DEAD box RNA helicases DDX5 and DDX17 in chromatin organization and transcriptional regulation. BMB Rep. 2018;51(12):613–22.

[30] Kwon HS, Koh SH. Neuroinflammation in neurodegenerative disorders: the roles of microglia and astrocytes. Transl Neurodegener. 2020;9(1):42.

[31] Sen MK, Mahns DA, Coorssen JR, Shortland PJ. The roles of microglia and astrocytes in phagocytosis and myelination: Insights from the cuprizone model of multiple sclerosis. Glia. 2022;70(7):1215–50.

[32] Colicelli J. Human RAS superfamily proteins and related GTPases. Sci STKE. 2004;2004(250):RE13.

[33] Reiner DJ, Lundquist EA. Small GTPases. WormBook. 2018;2018:1–65.

[34] Young LC, Rodriguez-Viciana P. MRAS: A Close but Understudied Member of the RAS Family. Cold Spring Harb Perspect Med. 2018;8(12):a033621.

[35] Young LC, Hartig N, Boned Del Río I, Sari S, Ringham-Terry B, Wainwright JR, et al. SHOC2-MRAS-PP1 complex positively regulates RAF activity and contributes to Noonan syndrome pathogenesis. Proc Natl Acad Sci U S A. 2018;115(45):E10576-85.

[36] Zhao A, Li D, Mao X, Yang M, Deng W, Hu W, et al. GNG2 acts as a tumor suppressor in breast cancer through stimulating MRAS signaling. Cell Death Dis. 2022;13(3):260.

[37] Song Y, Ma R, Zhang H. The influence of MRAS gene variants on ischemic stroke and serum lipid levels in Chinese Han population. Medicine (Baltimore). 2019;98(48):e18065.

[38] Craveri A, Tornaghi G, Paganardi L, Ranieri R, Di Bella M, Gallo E. Le alterazioni emoreologiche del globulo rosso in soggetti a rischio per aterosclerosi [Hemorrheologic disorders of red cells in subjects at risk for atherosclerosis]. Minerva Med. 1987;78(13):893–8.

[39] Mahmood F, Xu R, Awan MUN, Song Y, Han Q, Xia X, et al. PDIA3: Structure, functions and its potential role in viral infections. Biomed Pharmacother. 2021;143:112110.

[40] Lam STT, Lim CJ. Cancer Biology of the Endoplasmic Reticulum Lectin Chaperones Calreticulin, Calnexin and PDIA3/ERp57. Prog Mol Subcell Biol. 2021;59:181-96.

[41] Cassano T, Giamogante F, Calcagnini S, Romano A, Lavecchia AM, Inglese F, et al. PDIA3 Expression Is Altered in the Limbic Brain Regions of Triple-Transgenic Mouse Model of Alzheimer’s Disease. Int J Mol Sci. 2023;24(3):3005.

[42] Hasel P, Aisenberg WH, Bennett FC, Liddelow SA. Molecular and metabolic heterogeneity of astrocytes and microglia. Cell Metab. 2023;35(4):555–70.

[43] Donnelly CR, Andriessen AS, Chen G, Wang K, Jiang C, Maixner W, et al. Central Nervous System Targets: Glial Cell Mechanisms in Chronic Pain. Neurotherapeutics. 2020;17(3):846–60.

[44] Lee HG, Lee JH, Flausino LE, Quintana FJ. Neuroinflammation: An astrocyte perspective. Sci Transl Med. 2023;15(721):eadi7828.

[45] Prydz A, Stahl K, Zahl S, Skauli N, Skare Ø, Ottersen OP, et al. Pro-Inflammatory Role of AQP4 in Mice Subjected to Intrastriatal Injections of the Parkinsonogenic Toxin MPP. Cells. 2020;9(11):2418.

[46] Pisani F, Simone L, Mola MG, De Bellis M, Frigeri A, Nicchia GP, et al. Regulation of aquaporin-4 expression in the central nervous system investigated using M23-AQP4 null mouse. Glia. 2021;69(9):2235–51.

[47] Duan Y, Wang Y, Liu Y, Jin Z, Liu C, Yu X, et al. Circular RNAs in Parkinson’s Disease: Reliable Biological Markers and Targets for Rehabilitation. Mol Neurobiol. 2023;60(6):3261–76.

[48] Kalyanaraman B, Cheng G, Hardy M, You M. OXPHOS-targeting drugs in oncology: new perspectives. Expert Opin Ther Targets. 2023;27(10):939–952. doi:10.1080/14728222.2023.2261631

[49] D’Andrea G, Nordera G, Pizzolato G, Bolner A, Colavito D, Flaibani R, et al. Trace amine metabolism in Parkinson’s disease: low circulating levels of octopamine in early disease stages. Neurosci Lett. 2010;469(3):348–51.

[50] Cossu G, Melis M, Sarchioto M, Melis M, Melis M, Morelli M, et al. 6-n-propylthiouracil taste disruption and TAS2R38 nontasting form in Parkinson’s disease. Mov Disord. 2018;33(8):1331–9.

[51] Laurencin C, Timestit N, Marques A, et al. Efficacy and safety of clonidine for the treatment of impulse control disorder in Parkinson’s disease: a multicenter, parallel, randomised, double-blind, Phase 2b Clinical trial. J Neurol. 2023;270(10):4851-4859. doi:10.1007/s00415-023-11814-y

[52] Miyazono K. TGF-beta signaling by Smad proteins. Cytokine Growth Factor Rev. 2000;11(1-2):15–22.

[53] Hampel H, Caraci F, Cuello AC, Caruso G, Nisticò R, Corbo M, et al. A Path Toward Precision Medicine for Neuroinflammatory Mechanisms in Alzheimer’s Disease. Front Immunol. 2020;11:456.

[54] Bito H, Takemoto-Kimura S. Ca(2+)/CREB/CBP-dependent gene regulation: a shared mechanism critical in long-term synaptic plasticity and neuronal survival. Cell Calcium. 2003;34(4-5):425–30.

[55] Zhang L, Zhang Y, Wang C, Yang Y, Ni Y, Wang Z, et al. Integrated single-cell RNA sequencing analysis reveals distinct cellular and transcriptional modules associated with survival in lung cancer. Signal Transduct Target Ther. 2022;7(1):9.

[56] Huang J, Liu L, Qin L, Huang H, Li X. Single-Cell Transcriptomics Uncovers Cellular Heterogeneity, Mechanisms, and Therapeutic Targets for Parkinson’s Disease. Front Genet. 2022;13:686739.

[57] Mutez E, Larvor L, Leprêtre F, Mouroux V, Hamalek D, Kerckaert JP, et al. Transcriptional profile of Parkinson blood mononuclear cells with LRRK2 mutation. Neurobiol Aging. 2011;32(10):1839–48.

[58] Ji X, Huang C, Mao H, Zhang Z, Zhang X, Yue B, et al. Identification of immune- and autophagy-related genes and effective diagnostic biomarkers in endometriosis: a bioinformatics analysis. Ann Transl Med. 2022;10(24):1397.

[59] Luo L, Zhu J, Guo Y, Li C. Mitophagy and immune infiltration in vitiligo: evidence from bioinformatics analysis. Front Immunol. 2023;14:1164124.

[60] Butler A, Hoffman P, Smibert P, Papalexi E, Satija R. Integrating single-cell transcriptomic data across different conditions, technologies, and species. Nat Biotechnol. 2018;36(5):411–420.

[61] Satija R, Farrell JA, Gennert D, Schier AF, Regev A. Spatial reconstruction of single-cell gene expression data. Nat Biotechnol. 2015;33(5):495–502.

[62] Huang J, Liu L, Qin L, Huang H, Li X. Single-Cell Transcriptomics Uncovers Cellular Heterogeneity, Mechanisms, and Therapeutic Targets for Parkinson’s Disease. Front Genet. 2022;13:686739.

[63] Aran D, Looney AP, Liu L, Wu E, Fong V, Hsu A, et al. Reference-based analysis of lung single-cell sequencing reveals a transitional profibrotic macrophage. Nat Immunol. 2019;20(2):163–172.

[64] Jin S, Guerrero-Juarez CF, Zhang L, Chang I, Ramos R, Kuan CH, et al. Inference and analysis of cell-cell communication using CellChat. Nat Commun. 2021;12(1):1088.

[65] Ritchie ME, Phipson B, Wu D, Hu Y, Law CW, Shi W, et al. limma powers differential expression analyses for RNA-sequencing and microarray studies. Nucleic Acids Res. 2015;43(7):e47.

[66] Fan LY, Yang J, Liu RY, Kong Y, Guo GY, Xu YM. Integrating single-nucleus sequence profiling to reveal the transcriptional dynamics of Alzheimer’s disease, Parkinson’s disease, and multiple sclerosis. J Transl Med. 2023;21(1):649.

[67] Ziffels B, Stringhini M, Probst P, Fugmann T, Sturm T, Neri D. Antibody-Based Delivery of Cytokine Payloads to Carbonic Anhydrase IX Leads to Cancer Cures in Immunocompetent Tumor-Bearing Mice. Mol Cancer Ther. 2019;18(9):1544–54.1

[68] Wu T, Hu E, Xu S, Chen M, Guo P, Dai Z, et al. clusterProfiler 4.0: A universal enrichment tool for interpreting omics data. Innovation (Camb). 2021;2(3):100141.

[69] Hemani G, Zheng J, Elsworth B, Wade KH, Haberland V, Baird D, et al. The MR-Base platform supports systematic causal inference across the human phenome. Elife. 2018;7:e34408.

[70] Bowden J, Davey Smith G, Burgess S. Mendelian randomization with invalid instruments: effect estimation and bias detection through Egger regression. Int J Epidemiol. 2015;44(2):512–25.

[71] Bowden J, Davey Smith G, Haycock PC, Burgess S. Consistent Estimation in Mendelian Randomization with Some Invalid Instruments Using a Weighted Median Estimator. Genet Epidemiol. 2016;40(4):304–14.

[72] Burgess S, Scott RA, Timpson NJ, Davey Smith G, Thompson SG; EPIC-InterAct Consortium. Using published data in Mendelian randomization: a blueprint for efficient identification of causal risk factors. Eur J Epidemiol. 2015;30(7):543–52.

[73] Hartwig FP, Davey Smith G, Bowden J. Robust inference in summary data Mendelian randomization via the zero modal pleiotropy assumption. Int J Epidemiol. 2017;46(6):1985–1998. doi:10.1093/ije/dyx102

[74] Wu T, Hu E, Xu S, Chen M, Guo P, Dai Z, et al. clusterProfiler 4.0: A universal enrichment tool for interpreting omics data. Innovation (Camb). 2021;2(3):100141.

[75] Martirosyan A, Ansari R, Pestana F, Hebestreit K, Gasparyan H, Aleksanyan R, et al. Unravelling cell type-specific responses to Parkinson’s Disease at single cell resolution [published correction appears in Mol Neurodegener. 2024 Mar 25;19(1):28.

[76] Aran D, Looney AP, Liu L, Wu E, Fong V, Hsu A, et al. Reference-based analysis of lung single-cell sequencing reveals a transitional profibrotic macrophage. Nat Immunol. 2019;20(2):163–72.

[77] Trapnell C, Cacchiarelli D, Grimsby J, Pokharel P, Li S, Morse M, et al. The dynamics and regulators of cell fate decisions are revealed by pseudotemporal ordering of single cells. Nat Biotechnol. 2014;32(4):381–6.

[78] Aibar S, González-Blas CB, Moerman T, Huynh-Thu VA, Imrichova H, Hulselmans G, et al. SCENIC: single-cell regulatory network inference and clustering. Nat Methods. 2017;14(11):1083–6.

[79] Su G, Morris JH, Demchak B, Bader GD. Biological network exploration with Cytoscape 3. Curr Protoc Bioinformatics. 2014;47:8.13.1-8.13.24.

[80] Xin S, Fang W, Li J, Li D, Wang C, Huang Q, et al. Impact of STAT1 polymorphisms on crizotinib-induced hepatotoxicity in ALK-positive non-small cell lung cancer patients. J Cancer Res Clin Oncol. 2021;147(3):725–37.

[81] Wei JP, Wen W, Dai Y, Qin LX, Wen YQ, Duan DD, et al. Drinking water temperature affects cognitive function and progression of Alzheimer’s disease in a mouse model. Acta Pharmacol Sin. 2021;42(1):45–54.

[82] Yoshizaki K, Furuse T, Kimura R, Tucci V, Kaneda H, Wakana S, et al. Paternal Aging Affects Behavior in Pax6 Mutant Mice: A Gene/Environment Interaction in Understanding Neurodevelopmental Disorders. PLoS One. 2016;11(11):e0166665.

[83] Huang J, Zhang J, Wang F, Zhang B, Tang X. Comprehensive analysis of cuproptosis-related genes in immune infiltration and diagnosis in ulcerative colitis. Front Immunol. 2022;13:1008146.

[84] Zhang X, Chao P, Zhang L, Xu L, Cui X, Wang S, et al. Single-cell RNA and transcriptome sequencing profiles identify immune-associated key genes in the development of diabetic kidney disease. Front Immunol. 2023;14:1030198.

[85] Tang Q, Chen J, Di Z, Yuan W, Zhou Z, Liu Z, Han S, et al. TM4SF1 promotes EMT and cancer stemness via the Wnt/β-catenin/SOX2 pathway in colorectal cancer. J Exp Clin Cancer Res. 2020;39(1):232.

